# Anthropogenic supply of nutrients in a wildlife reserve may compromise conservation success

**DOI:** 10.1101/2022.09.20.508584

**Authors:** Andrew J Abraham, Ethan Duvall, Elizabeth le Roux, Andre Ganswindt, Marcus Clauss, Christopher Doughty, Andrea Webster

## Abstract

In nutrient-poor wildlife reserves it has become common-place to provide supplemental mineral resources for wildlife. Yet, the impacts of anthropogenic mineral supplementation on community-wide wildlife nutrition, behaviour and subsequent impact on ecosystem processes remain poorly understood. Here, we examine the contribution of anthropogenic mineral lick provision to wildlife nutrient intake across a community of large mammals (>10kg) in the southern Kalahari Desert. Based on predicted daily nutrient requirements and a faecal nutrient assessment, large herbivores appear deficient in phosphorus (P), sodium (Na) and zinc (Zn). For these nutrients, anthropogenic salt and mineral licks constitute an important (>10%) source of nutrient intake helping to reduce or overcome requirement deficits. Larger-bodied species disproportionately consumed licks (p<0.01), acquiring more nutritional benefits. A comprehensive assessment of animal body condition indicated that in general large herbivores display good health. However, bulk grazers, hindgut fermenters and females were more likely to display signs of malnourishment. We discuss how provisioning of anthropogenic mineral licks may be inflating large herbivore populations beyond the long-term carrying capacity of the reserve, with subsequent impacts for ecosystem integrity and herbivore population instability. Based on results presented here, it is clear that anthropogenic provision of mineral licks should be considered carefully by wildlife managers aiming to conserve or restore natural processes in conservation and rewilding landscapes.

## 1. Introduction

The nutritional status of wildlife directly influences animal health, fertility and susceptibility to disease and predation (Robbins, 1993; Milner et al., 2014). Consequently, human activities that limit wildlife access to nutrient resources can exert a strong control over ecosystem composition and function (Hobbs and Swift, 1985; Birnie-Gauvin et al., 2017; Abraham et al., 2022). For example, many protected areas (PAs) throughout the world are fenced to mitigate human-wildlife conflicts (Pekor et al., 2019), which prevents free movement of large-bodied animals and may limit access to essential resources such as food and water (Jakes et al., 2018). Particularly in oligotrophic environments, where these animals tend to have larger home ranges, fences can cause substantial nutritional stress (Boone and Hobbs, 2004).

To counter impeded access to adequate nutrition in fenced, oligotrophic PAs, wildlife managers often supply anthropogenic mineral licks (Milner et al., 2014; Murray et al., 2016; Simpson et al., 2020). The benefits of supplemental nutrition for animal health have been well understood in livestock and hunting landscapes for thousands of years (Theiler et al., 1924; Oro et al., 2013). In these settings, the aim is to maximise physical growth and population density of target species for increased harvest, protein or economic return (McDowell, 1996; Bartoskewitz et al., 2003). Similar motives exist in tourism landscapes, where increasing the health, density and encounter rate of wildlife via mineral provisioning can lead to enhanced viewing experiences for tourists (Dubois and Fraser, 2013; Cox and Gaston, 2018).

While supplementing minerals in PAs has become common practice, it has largely retained the production-oriented approach inherited from its agricultural origins. Yet, the impacts of anthropogenic lick provision on community-wide wildlife nutrition, behaviour and subsequent impact on ecosystem processes have received little attention (Brown and Cooper, 2006; Milner et al., 2014). For example, in PAs where the aim is to preserve or restore natural processes, what quantity of anthropogenic mineral lick is the correct amount to provide? On one hand, too little provision may result in malnourished individuals that are susceptible to disease or predation, raising questions of ethical animal husbandry (Dubois and Fraser, 2013). On the other, excessive provision may result in exceptionally healthy animals that inflate population densities above the long-term carrying capacity of the ecosystem (Oesterheld et al., 1992; Milner et al., 2014). In the latter case, this may compromise the integrity of the PA due to issues of over-grazing or herbivore population instability (Boone and Hobbs, 2004; Milner et al., 2014). Accordingly, if anthropogenic licks are provided in PAs, it is important to ascertain which species/individuals consume the licks, what contribution this makes to their required nutrient intake, and whether supplementation helps to reach conservation goals.

Here, we examine the contribution of anthropogenic mineral lick provision for wildlife nutrient intake across a large (>10kg) mammal community in the southern Kalahari Desert. Given this region is nutrient-poor (O’Halloran et al., 2010), mineral licks may form a critical part of nutrient intake for large vertebrate populations (Knight et al., 1988). Specifically, we aim to quantify the proportion of required nutrients that large mammalian herbivores acquire from anthropogenic salt and mineral licks compared to other primary sources of forage and water. We then assess what role these additional nutrients may have for wildlife health by undertaking two separate measures; (i) an assessment of faecal nutrient concentrations, and (ii) an assessment of animal body condition. Both measures have been widely employed for non-invasive assessments of wildlife health and can provide a holistic picture of nutrition status (Wrench et al., 1997; Ezenwa et al., 2009). Finally, we discuss the role that anthropogenic mineral provision may have for reaching conservation goals and how to ensure that wildlife management decisions are appropriate to reach these.

## 2. Methods

### 2.1 Study site

Tswalu Kalahari Reserve (TKR) is a 121,700 ha fenced wildlife reserve located at -27.26, 22.45 in the southern Kalahari Desert, South Africa (figure 1). Prior to 1995, TKR comprised ∼40 domestic livestock farms, but was converted to a wildlife reserve by the removal of internal fences and associated infrastructure. The foundational principle of TKR is ecological restoration financed by tourism (https://tswalu.com/conservation-story/conservation/). Today, TKR includes a complement of large vertebrate herbivores native to the region, as well as a number of species that would historically have occurred seasonally, but are now resident within the fenced system (see table 1; van Rooyen and van Rooyen, 2022). An electrified fence divides the reserve into two sections; the Greater Korannaberg section (92,231 ha), which harbours cheetah (*Acinonyx jubatus;* 4.3 ind.100 km^-2^), African wild dog (*Lycaon pictus*; 0.4 ind.100 km^-2^) and spotted hyaena (*Crocuta crocuta*; 1.3 ind.100 km^-2^), and the Lekgaba section (18,649 ha), which supports two prides of lion (*Panthera leo*; 5.9 ind.100 km^-2^) (figure 1).

**Figure 1.**
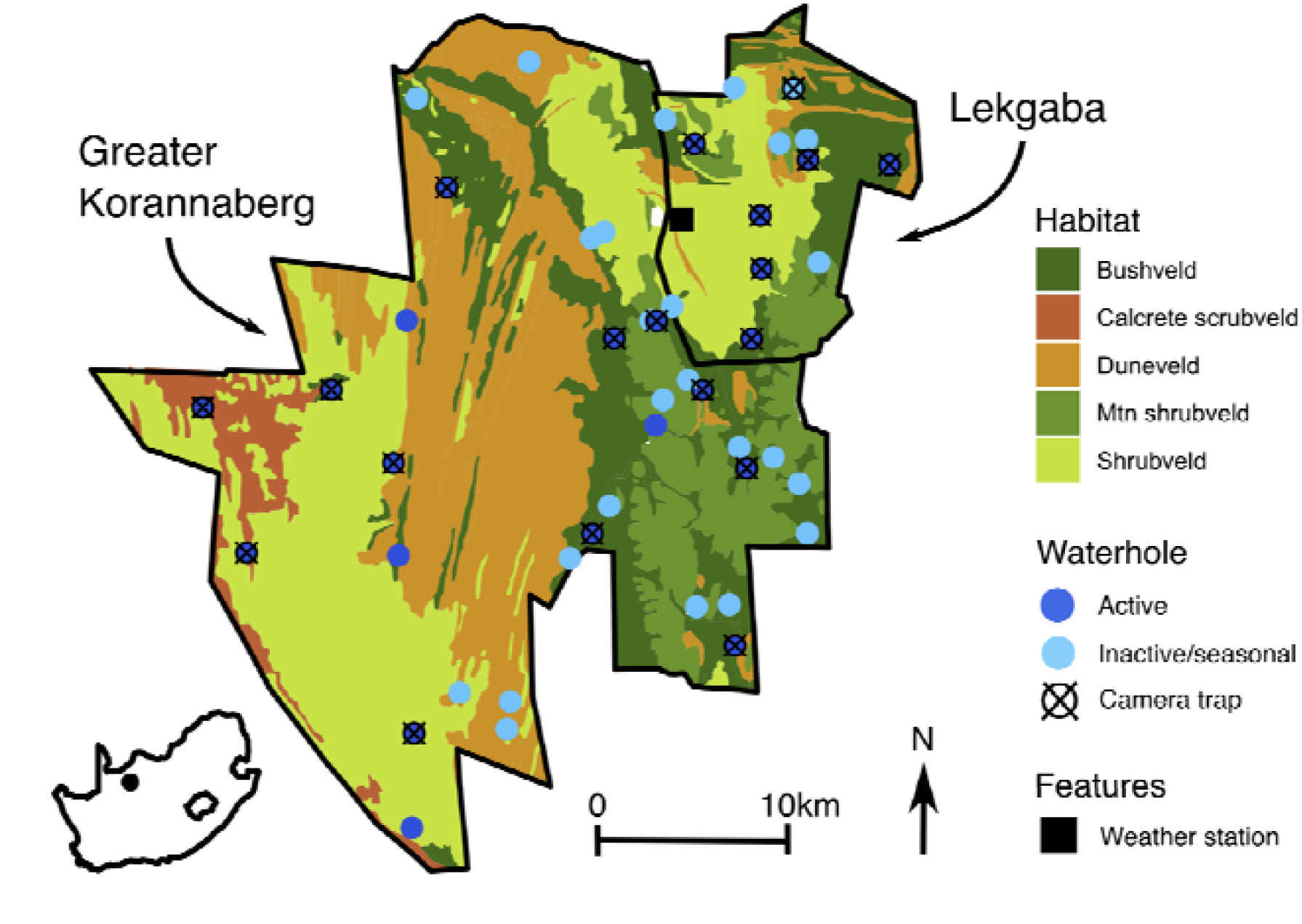
Tswalu Kalahari Reserve (TKR), South Africa, highlighting the Greater Korannaberg and Lekgaba sections, major habitat types (van Rooyen and van Rooyen, 2022) and the location of waterholes. Sites where camera traps were placed at salt and mineral licks to observe utilisation by large herbivores (>10kg) are denoted.

**Table 1.**
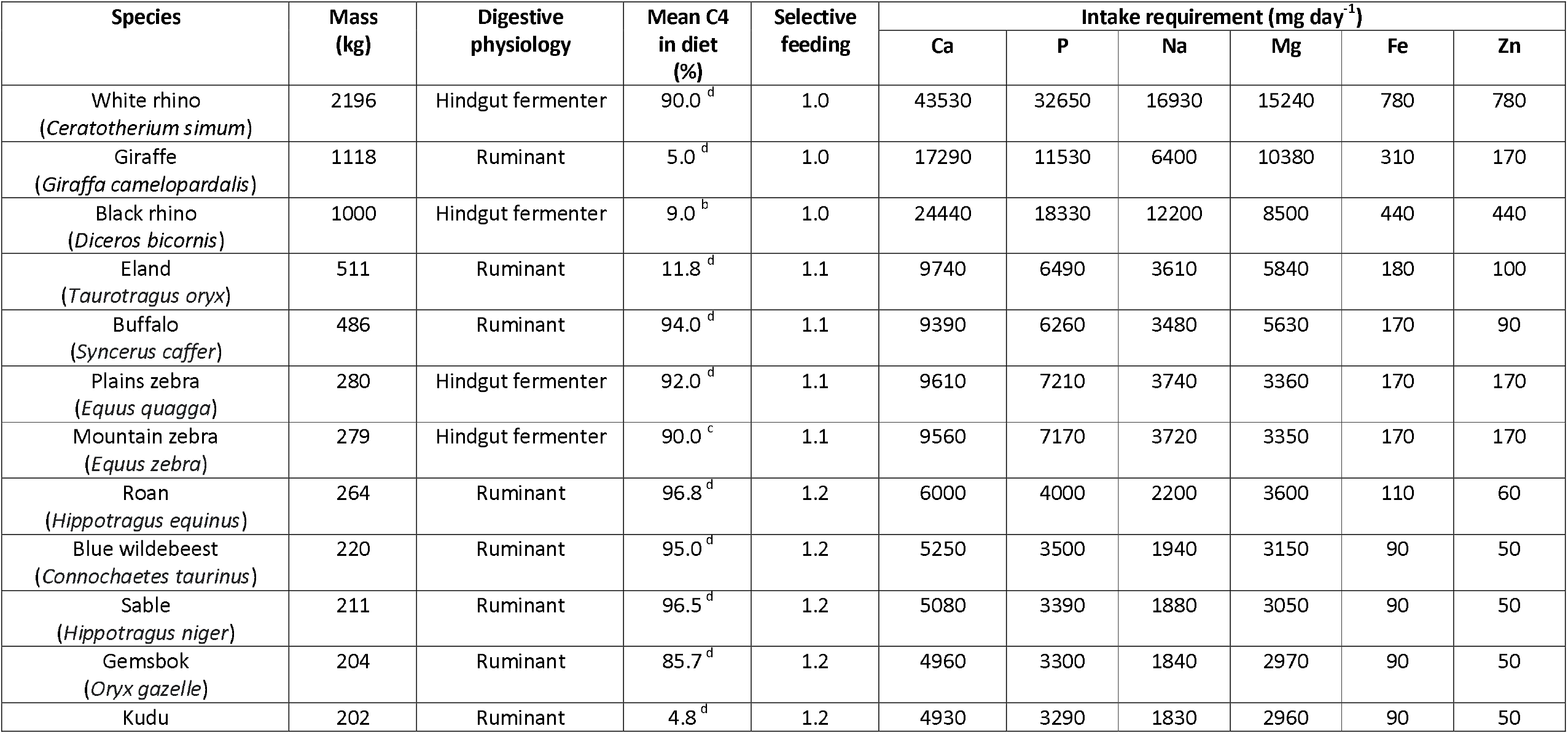
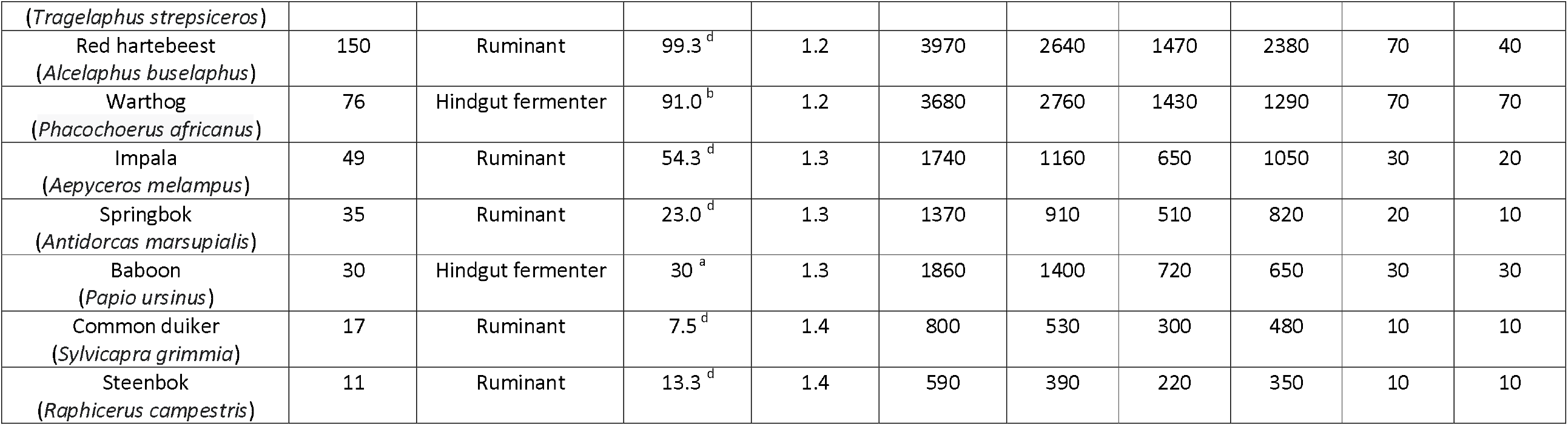
Characteristics and nutrient requirements for maintenance in ungulates resident at Tswalu Kalahari Reserve. Body mass values taken from Hempson et al. (2015), dietary C4 % from ^a^Codron et al. (2006), ^b^Codron et al. (2007), ^c^Strauss (2015) and ^d^Staver and Hempson (2020) and nutrient requirements from Lintzenich et al. (1997).

Much of TKR is underlain by aeolian sands of the Gordonia formation, with the emerging Korannaberg mountains formed of subgraywacke, quartzite, slate, dolomite, jasper and conglomerate (van Rooyen and van Rooyen, 2022). Sands of the southern Kalahari contain low concentrations of many critical nutrients for animal health, including nitrogen (N), phosphorus (P), sodium (Na), copper (Cu) and zinc (Zn) (Knight et al., 1988; Cromhout, 2007; O’Halloran et al., 2010). As a result, wildlife managers at TKR provide supplementary nutrients in the form of anthropogenic ∼62kg salt (Na) and ∼25kg mineral (Ca, P, S, Co, Cu, I, Fe, Mg, Se, Zn) licks distributed throughout the reserve (figure 2; Abraham et al., 2021). These licks are purchased from local suppliers (https://safarifeeds.co.za/products) and placed near water sources throughout TKR (Greater Korannaberg 15 sites, Lekgaba 7 sites; see figure 1). Once licks are consumed, they are immediately replenished ensuring constant availability throughout the year. In total, TKR provide 8-10,000 kg salt and 20-25,000 kg mineral lick per year across the reserve. Information pertaining to lick provision for the Greater Korannaberg and Lekgaba sections separately is not available. Consequently, in this paper both sections together are considered together.

**Figure 2.**
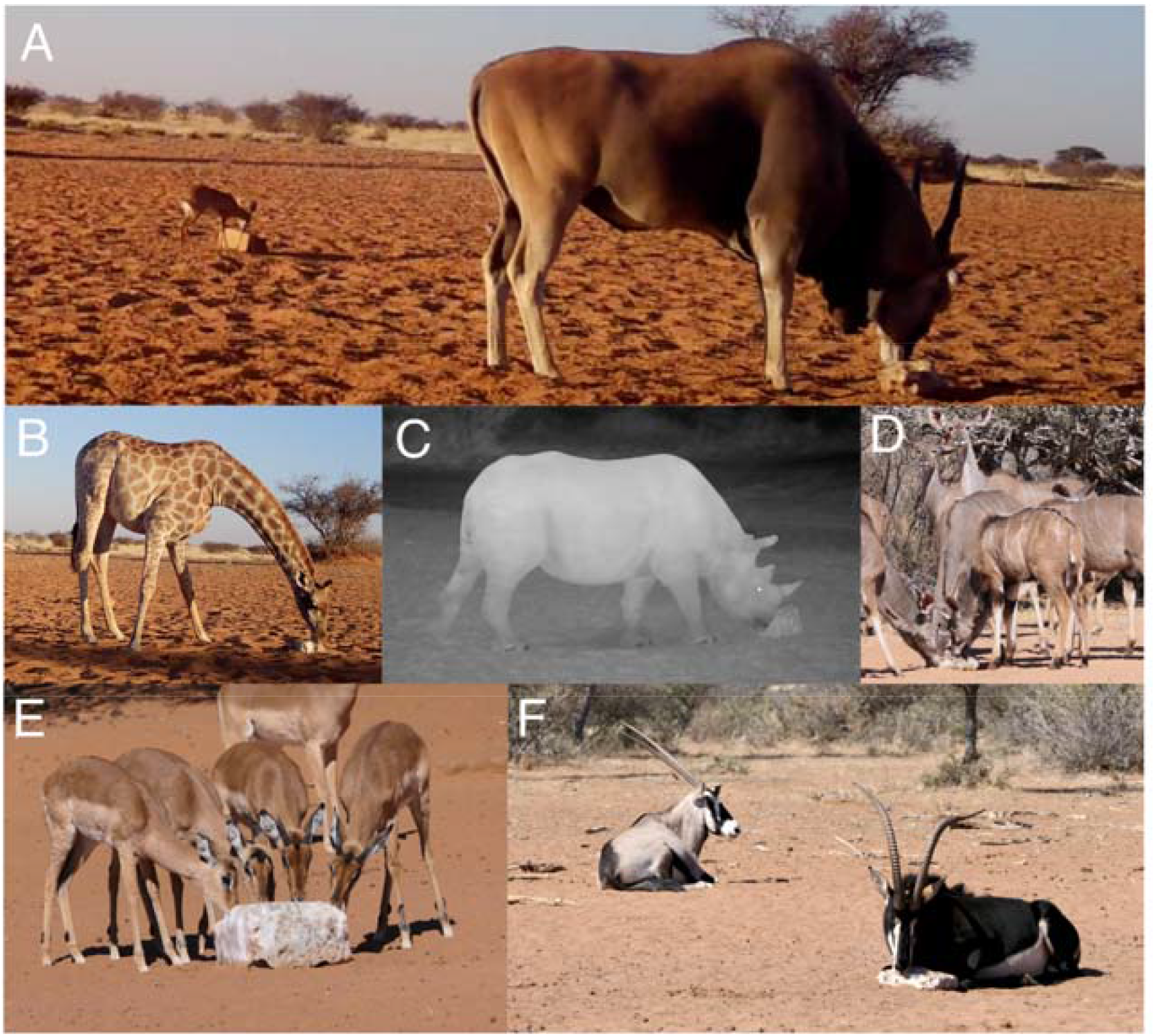
*Salt and mineral lick consumption by (A) steenbok (Raphicerus campestris*) and eland (*Taurotragus oryx*), (B) giraffe (*Giraffa Camelopardalis*), (C) black rhino (*Diceros bicornis*). (D) kudu (*Tragelaphus strepsiceros*), (E) impala (*Aepyceros melampus*), and (F) gemsbok (*Oryx gazelle*) and sable (*Hippotragus niger*) at Tswalu Kalahari Reserve (TKR). Photos taken by A. Abraham and A. Webster.

### 2.2 Large herbivore abundance

At TKR, large herbivore (>10kg) aerial counts are conducted annually during the beginning of the dry season (March-April). We transformed raw count data for each species into an estimate of abundance using the principles of ‘distance’ sampling (see SI text 1 for details and comparison to ground-based validation; Buckland et al., 1993). Large herbivore abundance estimates for the Greater Korannaberg and Lekgaba sections in 2021 are provided in SI table 1.

### 2.3 Large herbivore nutrient requirements

Nutrient requirements have been extensively studied in livestock (Suttle, 2010). However, less research has been conducted on wild ungulate species, which may have different nutrient demands (Robbins, 1993). In addition, specific requirements within a species may vary due to environmental (e.g. climate), reproductive state (e.g. pregnancy or lactation) or individual (e.g. age, sex, parasite load) differences (Robbins, 1993; Suttle, 2010). Here, we take a conservative approach and compare mean annual nutrient intake from forage, water and anthropogenic mineral licks at a population level to optimal requirements suggested by Lintzenich et al. (1997) for adult zoo animals at maintenance. This allows us to contextualise the contribution of anthropogenic mineral licks, but overlooks critical seasonal and individual differences. Importantly, however, Lintzenich et al. (1997) do include the influence of body size and digestive tract morphology on nutritional requirements, which potentially modify specific requirements in mammals (Clauss et al., 2013). For example, hindgut fermenters often require higher nutrient intake due to their lower digestive efficiency (Clauss et al., 2015). Here, we specifically focus on Ca, P, Na, Zn, iron (Fe) and magnesium (Mg), which are all nutrients closely linked to health and reproduction in large mammals (Robbins, 1993; Suttle, 2010). Species-specific nutrient requirements were calculated using equation 1:

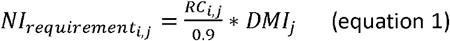

Where Ni_requirement,I,j_ is the required intake of nutrient *i* for species *j* in mg.day^-1^, RC is the required concentration of nutrient *i* for adult maintenance in mg.g dry matter^-1^ on a 90% dry matter basis for species *j* defined by Lintzenich et al. (1997), and DMI_j_ is dry matter intake estimated using the allometric relationship for field metabolic rate of 4.82*BM^0.734^ and metabolizable energy of the diet for species *j* equal to 10 kJ.g DM^-1^ for hindgut fermenters and 11.5 kJ.g DM^-1^ for ruminants from Nagy et al. (1999). Table 1 outlines species-specific mass, digestive physiology and daily nutrient intake requirements.

### 2.4 Large herbivore nutrient intake

In natural environments, large mammalian herbivores primarily acquire nutrients from forage and water (Robbins, 1993). Our aim here is to put nutrient acquisition from anthropogenic mineral licks at TKR into context when compared to other these major sources. Whilst geophagy (ingestion of soil) and osteophagy (ingestion of bone) are known contributors of nutrition for Kalahari herbivores (Knight et al., 1988; Holdo et al., 2002), we do not currently have estimates for these sources.

#### Natural forage intake

Plant nutrient content notably varies between grasses, forbs and woody plants (Kattage et al., 2021). Therefore to include diet differences, we first derived the contribution of C4 plants (i.e. grasses) to the diet of each species (see Table 1; Codron et al., 2007; Staver and Hempson, 2020). Nutrient concentrations for commonly-consumed plant species were then collated from studies previously conducted at TKR, representing seven C3 and six C4 species (Cromhout, 2007; Abraham et al., in prep, Webster et al., 2021; Prayag et al., 2020; Helary et al., 2012). Foliar concentration data from collated studies was, however, biased towards the dry season. In addition, the southern Kalahari experiences significant inter-annual climate variability (e.g. precipitation), which can influence nutritional quality (Skarpe and Bergstrom, 1987). To ensure that plant nutrient values reported at TKR are broadly representative of longer-term trends, we compared mean reported values at TKR to other sites in the southern Kalahari (table 2). In general, reported plant nutrient concentrations were similar to nearby sites for both C3 and C4 plants (table 2). We did, however, adjust excessively high reported values of Ca in C3 species at TKR from 30,765 mg.kg^-1^ to 15,000 mg.kg^-1^ to align with data reported from other sites. As herbivore diet quality generally scales negatively with body mass (Clauss et al., 2013), we modified plant nutrient concentration by a quality factor for each species to account for selective foraging by smaller species (see table 1). Daily forage nutrient intake for an individual of each species was calculated using equations 2 and 3:

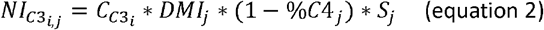

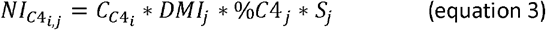

**Table 2.** Mean values [range in brackets] of nutrient concentrations from sources at Tswalu Kalahari Reserve. For comparison, forage nutrient concentrations in the southern Kalahari are provided from Skarpe and Bergstrom (1987), Tolsma et al. (1987), Stepelberg et al. (2008) and Theart (2015). Values from Tolsma et al. (1987) were extracted using WebPlotDigitizer (https://apps.automeris.io/wpd/).

| Source | Location | Ca<br>(mg kg<sup>-1</sup>) | P<br>(mg kg<sup>-1</sup>) | Na<br>(mg kg<sup>-1</sup>) | Mg<br>(mg kg<sup>-1</sup>) | Fe<br>(mg kg<sup>-1</sup>) | Zn<br>(mg kg<sup>-1</sup>) |
| --- | --- | --- | --- | --- | --- | --- | --- |
| C4 plants | Tswalu | 3727<br>[2000-7600] | 568<br>[300-1000] | 345<br>[164-680] | 2454<br>[500-11200] | 194<br>[83-475] | 16<br>[5-35] |
|  | Skarpe and Bergstrom (1987) | 3406<br>[2900-3956] | 768<br>[571-975] |  |  |  |  |
|  | Tolsma et al. (1987) | 8733<br>[2237-19635] | 956<br>[123-2543] |  | 2671<br>[425-6450] | 440<br>79-1062] |  |
|  | Stepelberg et al. (2008) | 6874<br>[6233-7213] | 900<br>[567-1350] |  |  |  |  |
| C3 plants | Tswalu | 30765<br>[29300-32230] | 1460<br>[1440-1480] | 176<br>[85-400] |  | 301<br>[175-530] | 14<br>[3-25] |
|  | Skarpe and Bergstrom (1987) | 12235<br>[7667-24000] | 1308<br>[550-2500] |  |  |  |  |
|  | Tolsma et al. (1987) | 16160<br>[8056-44850] | 1607<br>[712-2454] | 115<br>[11-193] | 2347<br>[1118-4155] | 167<br>[56-391] | 24<br>[13-52] |
|  | Stepelberg et al. (2008) | 15812<br>[9900-22950] | 1378<br>[825-1967] |  |  |  |  |
|  | Theart (2015) | 16450<br>[7450-26480] | 1170<br>[750-1870] |  |  |  | 20<br>[13-47] |
| Water | Tswalu | 38<br>[5-150] | 0.17<br>[0-1.67] | 24<br>[6-70] | 19<br>[3-91] | 0.06<br>[0-0.7] | 0<br>[0-0.0001] |
| Anthropogenic salt lick | Tswalu | 4133<br>[4025-4194] | 74<br>[13-178] | 191979<br>[188319-196499] | 143<br>[74-200] | 103<br>[46-200] | 0.7<br>[0.2-1.1] |
| Anthropogenic mineral lick | Tswalu | 83262<br>[77000-86897] | 36132<br>[29869-44291] | 55284<br>[46159-60635] | 3287<br>[2867-3794] | 1370<br>[1336-1398] | 181<br>[152-200] |

Where NI_C3,i,j_ and NI_C4,i,j_ is the nutrient intake from C3 and C4 plants of nutrient *i* for an individual of species *j* in mg.day^-1^, C_C3,i_ and C_C4,i_ are the average concentrations of nutrient *i* in mg.g dry matter^-1^ of C3 and C4 plants, DMI_j_ is dry matter intake for species *j* in g.day^-1^ (see equation 1), %C4_j_ is the percentage of C4 plants in the diet of species *j* and S_j_ is a forage selectivity factor for species *j* (ranging from highly selective, 1.4, to unselective, 1.0 – see table 1).

#### Water

Waterhole quality surveys have been conducted extensively at TKR over the period 2000-2021 (see SI text 2 for details). Here, we calculated the mean nutrient concentration from all waterholes sampled across TKR. Daily water nutrient intake for an individual of each species was estimated using the allometric relationship of daily water requirements from Calder and Braun (1983):

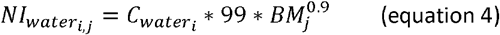

Where NI_Water,I,j_ is the nutrient intake from water of nutrient *i* for an individual of species *j* in mg.day^-1^, C_Water,I_ is the concentration of nutrient *i* in water in mg.litre^-1^, and BM is the body mass in kg for an individual of species *j*. Desert-adapted animals, however, often satisfy water consumption requirements through their diet rather than surface water (Kihwele et al., 2020). Consequently, the calculation of nutrient intake from water sources for herbivores at TKR is likely a conservative upper estimate.

#### Salt and mineral licks

To quantify nutrient intake from salt and mineral licks, we first analysed licks for nutrient concentration. 20g samples of salt (n=5) and mineral (n=5) licks were collected from TKR and individually pulverised. At the University of Pretoria’s Soil Sciences Laboratory, 0.25-0.30g of each sample was weighed into microwave digestion tubes. 10 ml of Suprapur Nitric acid (65%) was added to each sample and samples were left to digest overnight prior to analysis for Ca, P, Na, Mg, Fe, Zn concentration using a SPECRO GENESIS Inductively Coupled Plasma Optical Emission Spectrometer (ICP-OES). To quantify salt and mineral lick consumption by large herbivore species, Bushnell TrophyCam HD and Browning Recon Force ADVANTAGE camera traps were placed at lick sites for in the Greater Korannaberg (n=12) and Lekgaba (n=7) sections between July-September 2021. Camera traps were programmed to record 30s videos with a 5-minute interval between recordings and left for 14 days. We recorded all large herbivore species (>10kg) present within the camera trap frame and the number of animals that physically made use of the salt and mineral licks. To ensure each camera trap contributed equally to data analysis, visitation rate from each camera trap was standardised /24hrs. Based on camera trap videos, we assume that 80% of licks are consumed by large herbivores, with the rest being consumed by other fauna (e.g. birds) or lost to the environment. We cannot measure how much lick is consumed during each discrete feeding event, but we do know the total quantity consumed in TKR annually (∼9000 kg salt lick; ∼22,500 kg mineral lick). We therefore divide the total lick consumption over the course of a year by the abundance and proportional use of licks by each species, with the assumption that an individual consumes proportionally to their metabolic rate (i.e. BM^0.75^) during each feeding event. We calculated individual daily nutrient intake from salt and mineral lick consumption using equations 5 and 6:

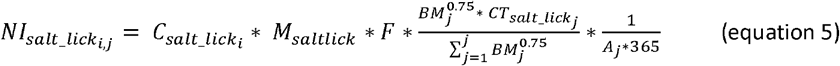

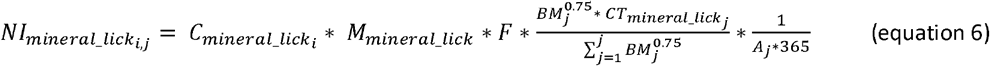

Where NI_salt_lick,i,j_ and NI_mineral_lick,i,j_ are the nutrient intake of nutrient *i* from salt lick and mineral licks respectively for an individual of species *j* in mg.day^-1^, C and C are the concentrations of nutrient *i* in salt and mineral licks in mg.g^-1^, M and M are the mass of licks distributed within TKR in g.year^-1^, F is the fraction of available lick consumed by large herbivores (>10kg) set to 0.8 (see above), BM_j_ is the body mass of species *j* (table 1), CT_salt_lick,j_ and CT_mineral_lick,j_ are the proportional use of salt and mineral licks by species *j* as determined by camera traps and A_j_ is the abundance of species *j* as determined by aerial counts.

### 2.5 Index of large herbivore nutrient stress

Acquiring direct measurements on the nutrient status of large wild herbivores is difficult without substantial and costly interference to the animals (Robbins, 1993; Birnie-Gauvin et al., 2017). Faecal nutrient analysis, however, offers a practical, non-invasive approach to measure nutrient stress and has been extensively applied to large herbivore species in southern Africa, including buffalo, zebra, giraffe, roan, kudu, springbok and elephant (e.g. Wrench et al., 1997; Grant et al., 2000; van der Waal et al., 2003; Stapelberg et al., 2008; Okita-Ouma et al., 2021). Faecal measurements reflect the quality and digestibility of resources consumed by an individual, and can be used to assess if the animal’s diet contains enough energy and protein to utilise additional nutrients acquired from mineral licks (McDowell, 1996; Steuer et al., 2014). If energy or protein are insufficient, the provision of additional nutrients serves no purpose and can even have negative effects for animal health by creating harmful free radicals leading to oxidative stress (Van Niekerk and Jacobs, 1985; Goff, 2018).

We collected fresh faeces (<10 minutes post-defecation) from three abundant ruminant species representing three feeding types; blue wildebeest (*Connochaetes taurinus*; obligate grazer), springbok (*Antidorcas marsupialis*; mixed feeder) and kudu (*Tragelaphus strepsiceros*; obligate browser). Faecal collection (60 per species; total n =180) was conducted coincident with camera trapping (July-September), when herbivores in the southern Kalahari are typically nutritionally stressed (Abraham et al., 2021). All faecal samples were lyophilised and homogenised prior to quantification of crude protein content, expressed as N, using a LECO® instrument equipped with TruMac CN/N determinator software (v1.5x) for instrument control and data processing. We then used acid digestion and ICP-OES quantification described above to determine P, Ca, Na, Fe, Mg and Zn concentration in the same herbivore faecal samples. We compared faecal macro-nutrient (N and P) concentrations to thresholds of potential concern (cut-off values below which mammalian herbivores have been documented suffering from growth and reproductive issues). For N we used 14 g.kg^-1^ (grazers; Wrench et al., 1997) and 15 g.kg^-1^ (mixed feeders and browsers; van der Waal et al., 2003) and for P we used 2 g.kg^-1^ (all species; Wrench et al., 1997). Different N thresholds were used for grazers and browsers due to the precipitating effect of condensed tannins elevating faecal N concentrations (Leslie et al., 2008; Steuer et al., 2014). Similar thresholds of potential concern have not been established for micro-nutrients and thus we focus our faecal analysis on N and P only.

### 2.6 Large herbivore body condition

Nutrition status has been directly linked to visual body condition in southern African herbivores (Grant et al., 2000; van der Waal et al., 2003; Lane et al., 2014). We therefore undertook a comprehensive assessment of large herbivore body condition using a visual assessment scheme that relies on objective assessment of clearly detectable characteristics (such as the protrusion of ribs vs the presence of fat deposits; see SI text 3 and SI table 2 for method for details; Ezenwa et al., 2009). Assessment was undertaken by an observer (AW) with >10 years professional experience observing wildlife in southern Africa. Body condition score (BCS; 1 low to 5 high), species and sex were recorded for all individuals encountered along 1009 km of driven road transects conducted coincident with the camera trap survey and faecal collection. If large herbivores are nutritionally compromised at TKR, low body condition scores would be expected.

### 2.7 Ethical statement

All data was collected with the approval of the University of Pretoria Research and Animal Use and Care Committee (Reference NAS115/2021) and the South African Department of Agriculture, Land Reform and Rural Development 12/11/1/1/8(1933NC) and 12/11/1/1/8 A (1933NC) JD.

## 3. Results

### 3.1 Contribution of anthropogenic mineral licks to herbivore nutrient requirements at TKR

From camera traps deployed at 19 anthropogenic mineral lick sites, we recorded 38,984 large vertebrate animal sightings. Specifically, 12,764 animals were recorded directly using salt licks and 5,722 utilising mineral licks. Lick site visits, salt lick use and mineral lick use relative to species abundance scaled positively with body mass (figure 3). This relationship was consistent between both the Greater Korannaberg and Lekgaba sections (SI figure 3). Buffalo (*Syncerus caffer*) relatively consumed salt licks the most (4.9x more regularly than predicted by their abundance), whilst plains zebra (*Equus quagga*) relatively consumed mineral licks the most (4.5x more regularly). Steenbok (*Raphicerus campestris*) very rarely used either salt or mineral licks, despite their high abundance at TKR (figure 3).

**Figure 3.**
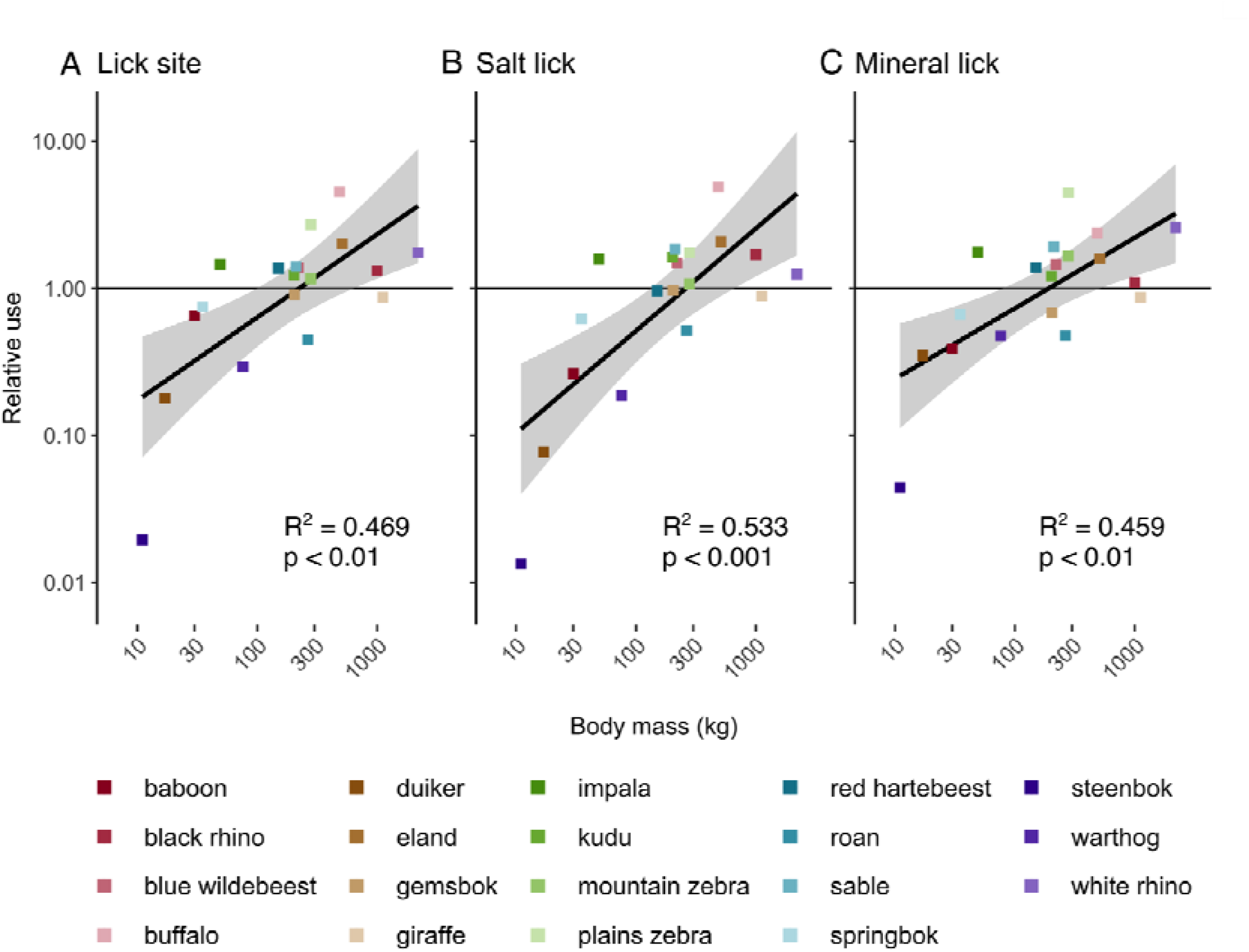
Relative visitation of large herbivores to anthropogenic lick sites (A), and direct use of salt (B) and mineral (C) licks compared to abundance at Tswalu Kalahari Reserve (TKR). Note that the y-axis is scaled log10, where values >1 represent more visitations of licks than expected based on species abundance and values <1 represent fewer visitations that expected. Trendlines represent a generalised-least squares model fit for all herbivores.

We calculated that many large herbivores at TKR consumed P, Na and Zn in amounts lower than required (figure 4). In general, grazers tended to be deficient in P, browsers deficient in Na and hindgut fermenters deficient in Zn. Salt and mineral licks constitute a substantial (>10%) component of daily intake of Na for all large herbivore species and of P for most large herbivores >100kg. Licks contributed >50% of daily Na intake for buffalo and eland (*Taurotragus oryx*) and >15% of daily P intake for buffalo, plains zebra and white rhino (*Ceratotherium simum*).

**Figure 4.**
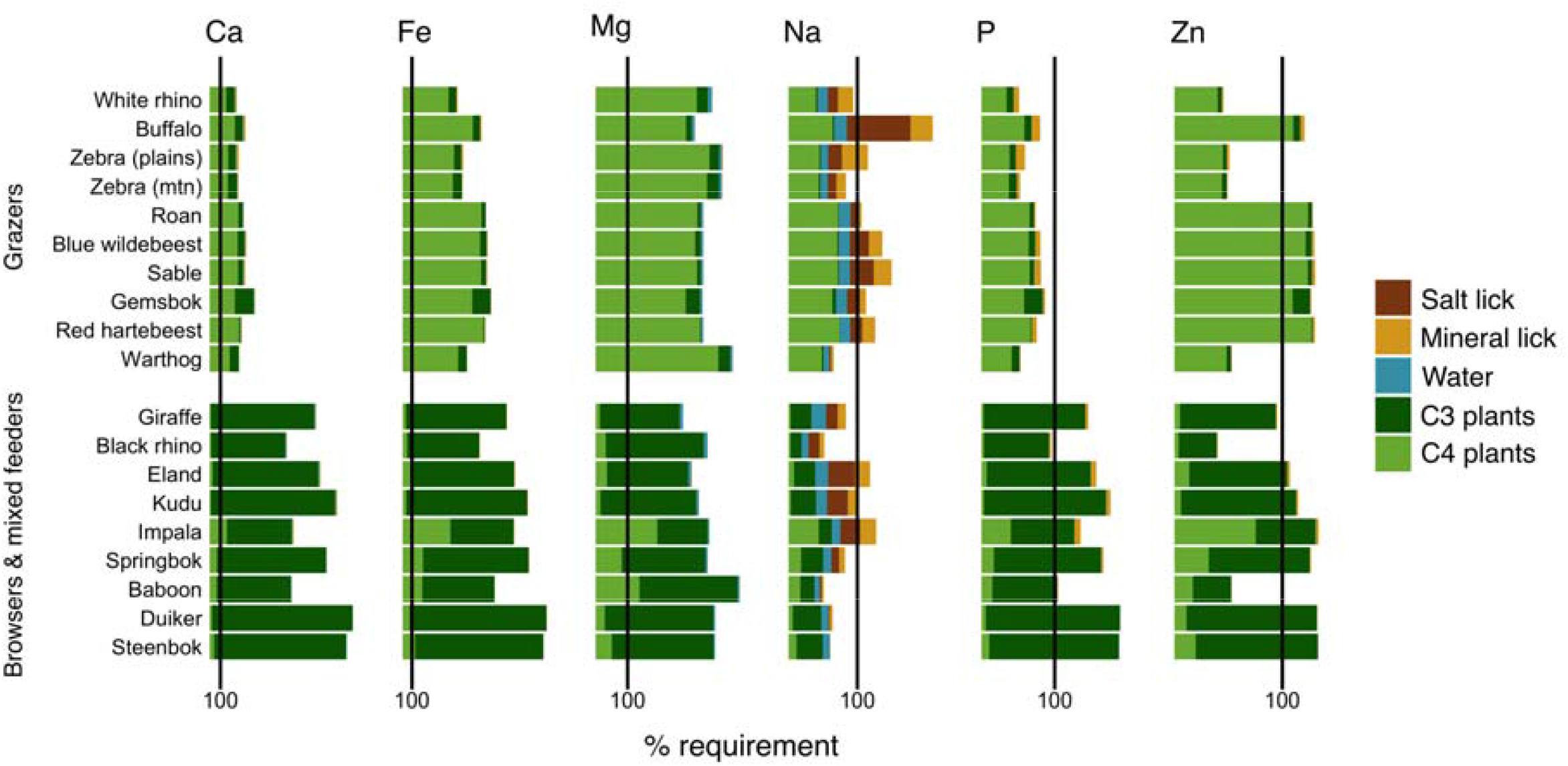
Daily intake of calcium (Ca), iron (Fe), magnesium (Mg), sodium (Na), phosphorus (P) and zinc (Zn) from different nutrient pools by large herbivores (>10kg) at Tswalu Kalahari Reserve (TKR). Vertical lines represent daily nutrient requirements for adult individuals at maintenance as suggested by Lintzenich et al. (1997).

### 3.2 Assessment of large herbivore health at TKR

92% of kudu (browsers) and 100% of springbok (mixed feeders) faecal samples are above the N threshold of potential concern, suggesting that most individuals from these groups may have sufficient protein in their diet to utilise provisional nutrients acquired from anthropogenic salt and mineral licks (figure 5). However, only 18% of blue wildebeest (grazer) faecal samples are above the critical N threshold of potential concern. Mean faecal P concentration is below the threshold of potential concern for blue wildebeest and kudu, whilst 43% of springbok samples are below the threshold of potential concern.

**Figure 5.**
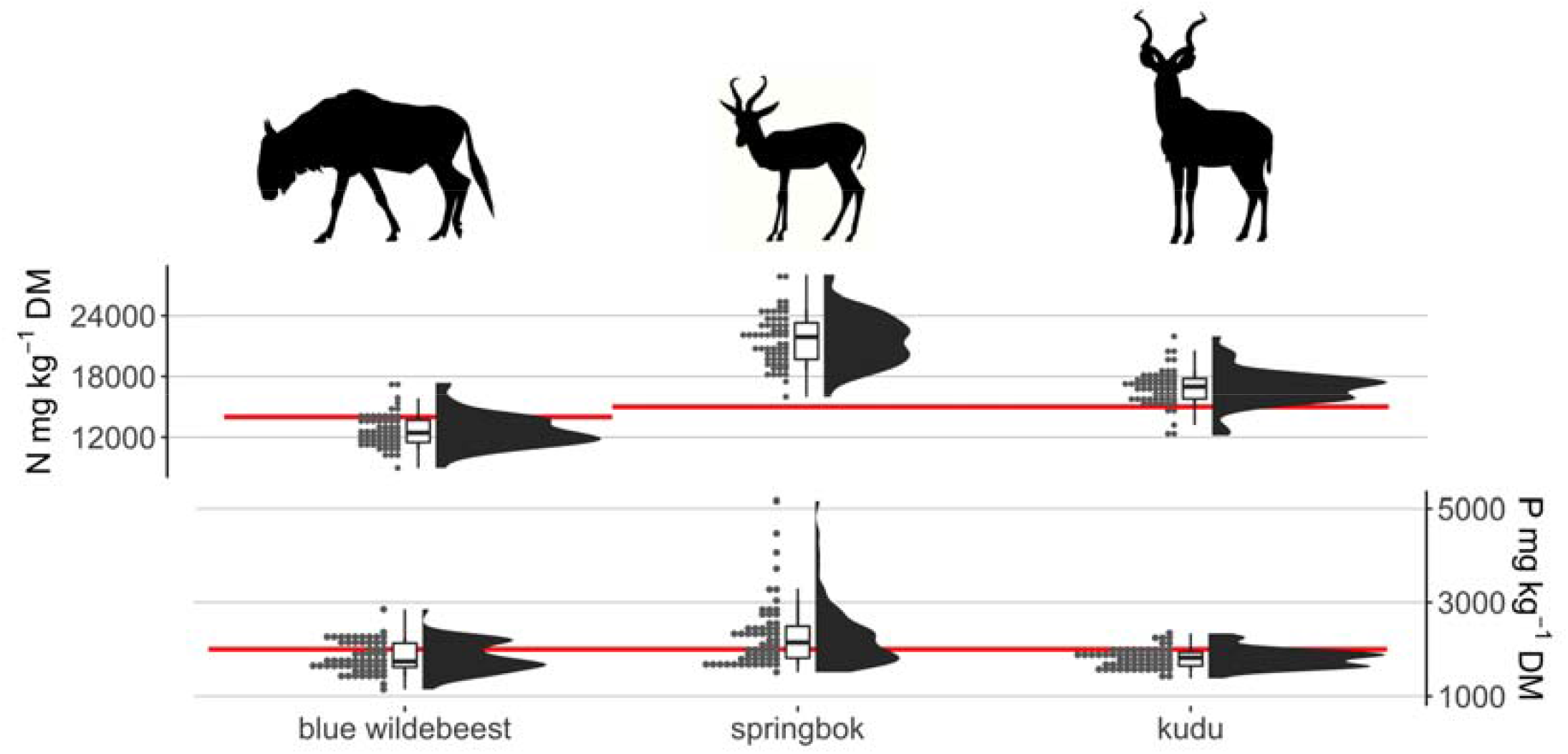
Faecal macro-nutrient concentrations of nitrogen (N) and phosphorus (P) for blue wildebeest (*Connochaetes taurinus*; obligate grazer), springbok (*Antidorcas marsupialis*; mixed feeder) and kudu (*Tragelaphus strepsiceros*; obligate browser) resident at Tswalu Kalahari Reserve (TKR). Red lines represent minimum faecal nutrient thresholds below which large mammalian herbivores begin suffering growth and reproductive issues. The N threshold is 14,000 mg kg^−1^ for grazers and 15,000 mg kg^−1^ for mixed feeders and browsers. The P threshold is 2,000 mg kg^−1^ for all species. Note that faeces were collected in the dry season (July-September) when nutrient stress is typically greatest.

In total, 1865 BCS were obtained from TKR. Generally, large herbivores (>10kg) are in reasonable health with a mean BCS of 2.97. Using two sample t-tests, grazers (M = 2.92; SD = 0.31) display a statistically significant lower body condition than mixed feeders and browsers M = 3.02; SD = 0.29), t(1744) = 6.879, p<0.001 (figure 6a), hindgut fermenters (M = 2.68; SD = 0.39) display a statistically significant lower body condition that ruminants (M = 2.99; SD = 0.28), t(165) = -9.51, p<0.001 (figure 6b), and females display a statistically significant lower body condition (M = 2.93; SD = 0.28) than males (M = 3.05; SD = 0.33), t(1050) = -6.36, p<0.001 (figure 6c).

**Figure 6.**
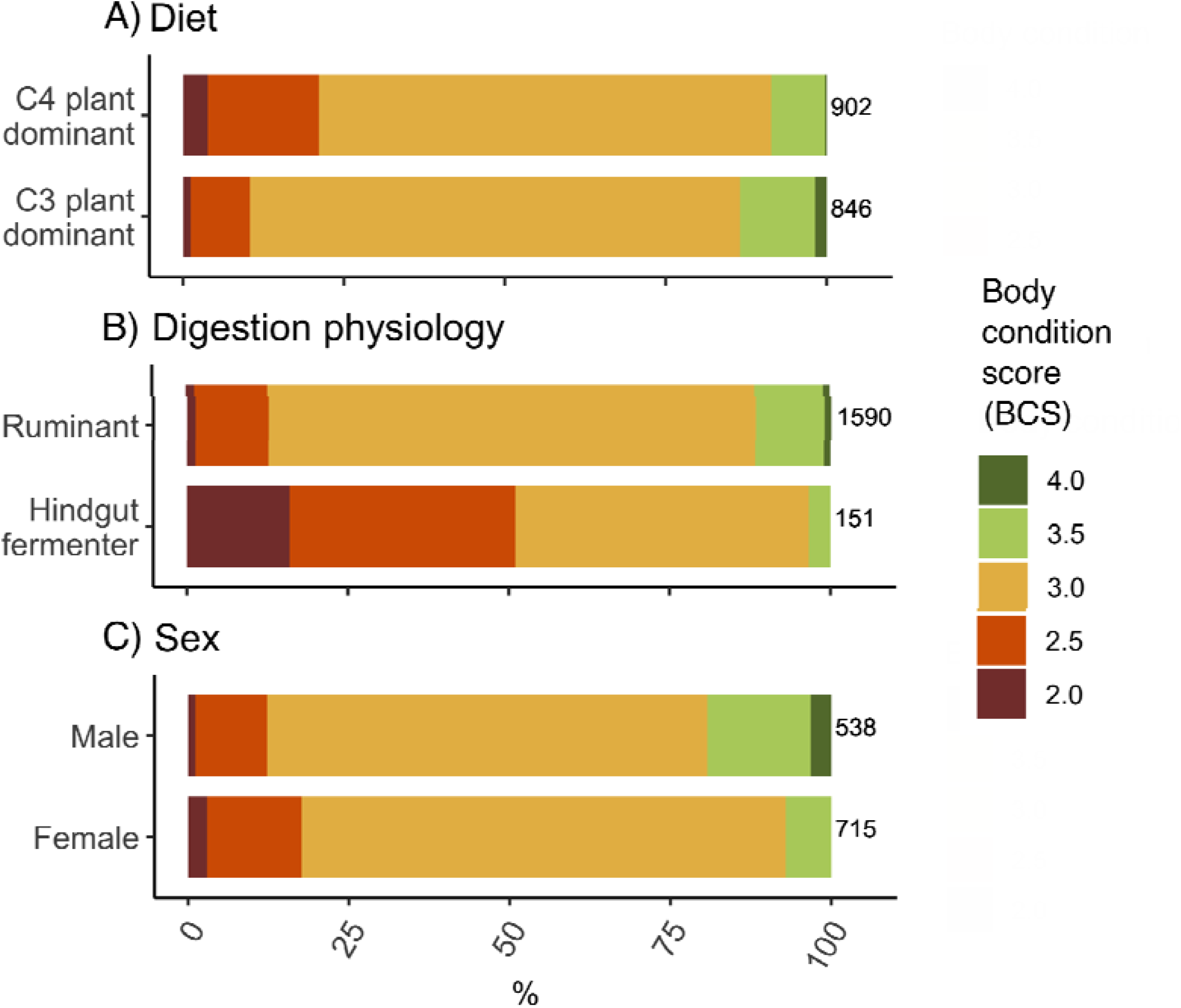
Body condition scores for large herbivore (>10kg) species at Tswalu Kalahari Reserve (TKR) split by diet, digestive physiology and sex groups. Numbers next to each bar denote sample size for that group.

## 4. Discussion

### 4.1 Mineral licks as an important source of nutrients at TKR

Mean daily salt lick consumption varies from 0.003 g.day^-1^ (steenbok) to 16.7 g.day^-1^ (buffalo), whilst mineral lick consumption varies from 0.02 g.day^-1^ (steenbok) to 68.4 g.day^-1^ (white rhino). These values compare similarly to the rates of anthropogenic lick consumption in other non-pastoral systems (e.g. ∼18 g.day^-1^ by white-tailed deer (*Odocoileus virginianus*) in Louisiana, USA (Schultz and Johnson, 1992). However, values are considerably lower than for livestock farming in low-nutrient environments, where lick site density is generally higher and predators are absent (e.g. 135±55 g cattle^-1^ day^-1^; Tait and Fisher, 1996).

Based on the theoretical daily nutrient requirement calculations and the faecal nutrient assessment conducted in this study, many large herbivores (>10kg) at TKR appear deficient in P, Na and Zn. For these nutrients, anthropogenic salt and mineral licks constitute an important (>10%) source of nutrient intake helping to reduce or overcome requirement deficits (figure 4). This is especially true for larger-bodied species that consume licks disproportionately (figure 3) and individuals that have elevated nutrient demands (e.g. pregnancy, lactation, juvenile; Suttle, 2010). It is important to note that the data used to estimate mineral requirements for this study (i.e. Lintzenich et al., 1997) do not represent determined minimum thresholds below which animal health is immediately compromised. Instead, they are descriptions from zoo nutritionists that contain an unknown safety margin, below which an individual’s health becomes suboptimal. Additionally, our calculations do not include other possible sources of nutrients for large herbivores at TKR including geophagy from natural mineral licks (Knight et al., 1988) and osteophagy (Bredin et al., 2008). If consumed at high rates, these alternative sources may offset the importance of anthropogenic mineral licks for species to acquire sufficient nutrient intake. Whilst geophagy is rarely seen at TKR, osteophagy is more commonly observed for several species including giraffe (*Giraffa camelopardalis*), sable (*Hippotragus niger*) and kudu (*Tragelaphus strepsiceros*) (Dylan Smith, Director of Research at TKR, personal communication). Bones, which have high P concentrations, may therefore provide an additional important P resource, especially for species with large skeletons, which increases P demand (Bredin et al., 2008).

Liebig’s Law of the Minimum suggests that anthropogenic nutrient supplementation is only important for large herbivores if energy and protein requirements are adequate (McDowell, 1996). Without sufficient calories, large herbivores cannot utilise supplementary nutrients for metabolic functions, growth and reproduction (Robbins, 1993). In some cases, additional nutrient intake from anthropogenic sources without sufficient energy and protein can even be harmful to wildlife. For example, in a factorial study examining the effect of energy, protein and P supplements on cattle fed low-quality forage in southern Africa, van Niekerk and Jacobs (1985) found that P supplementation had a negative effect on feed intake and body mass change, unless it was given in combination with both energy and protein supplements due to further burdening effects of P on an already unbalanced diet. Based on the faecal assessment for N conducted in this study, it appears that large herbivore grazers may not be able to utilise minerals from licks during the dry season due to a lack of dietary energy and protein. The mean dry season faecal N concentration for blue wildebeest of 12.6 g kg^-1^ falls below the threshold of potential concern equal to 14 g kg^-1^ as suggested by Wrench et al. (1997) (figure 5). In a focused study assessing the health of buffalo at TKR, Cromhout (2007) similarly found mean dry season faecal N concentrations below the threshold of potential concern at 10.8 g kg^-1^. Values this low indicate that some grazer individuals may struggle to maintain rumen fermentation due to inadequate nutrition for rumen microbes (Grant et al., 2000). However, in the same study, Cromhout (2007) also demonstrates that the N concentration of palatable grass species at TKR increases by 42-82% in the wet season, indicating that large herbivore grazers likely have sufficient energy/protein during this period to benefit from anthropogenic licks. Based on the results presented here, the majority of mixed feeders and browsers will, however, likely benefit from anthropogenic lick supplements throughout the year. The higher body condition scores of mixed feeders and browsers compared to grazers corroborates this assertion (figure 6a).

Where animals have sufficient energy and protein, many studies have highlighted the role of supplementary nutrients for rapidly increasing weight gain and increasing reproductive success in domestic (Tait and Fisher, 1996; de Brouwer et al., 2000) and wild animals (van der Waal et al., 2003; Milner et al., 2014). For example, a five-year study in the South African highveld found that significant P and Na supplementation of beef cattle during the wet summer season resulted in 18-27% higher body mass, 75-144% higher body condition score and 15-175% higher bone P content (de Brouwer et al., 2000). Consequently, the substantial contribution of salt and mineral licks to daily P and Na intake found in this study may play an important role for the health of large herbivores at TKR, albeit in times of the year when sufficient energy can be obtained. For grazers, anthropogenic licks may help individuals quickly build sufficient body condition during the wet season to buffer catabolism during the dry season (McDowell, 1996), whilst for mixed feeders and browsers licks may provide an important source of nutrients throughout the year (van der Waal et al., 2003). Although mineral licks contribute a smaller daily percentage of Zn (0.1-5.2%), this may be a critical source for hindgut fermenters, which have elevated demands due to antagonism of Zn with phytate reducing absorption efficiency for this group (Suttle, 2010). Zn-deficient animals tend to lose their appetite, which may help explain the considerably lower dry-season body condition of hindgut fermenters compared to ruminants (figure 6b).

### 4.2 Ecological implications of salt and mineral licks for conservation at TKR

Supplemental provision of salt and mineral licks at TKR has several ecological implications. First, the geography of anthropogenic lick sites strongly influences the movement and congregation patterns of many large herbivore species, with concomitant implications for species interactions, coexistence and localised sites of ecosystem impact (Priesmeyer et al., 2012). Second, nutrients gained from licks may have a strong influence on wildlife health, fertility and susceptibility to disease and predation (Suttle, 2010; Murray et al., 2016). In this study, we find that large mammalian herbivores generally display good body condition, despite our intake calculations and dry-season faecal assessments indicating deficiencies in P, Na and Zn (figures 4 and 5). Evidently, while these nutrients are low, they do not appear critically limiting for animal health. It is likely that the provision of salt and mineral licks, in combination with year-round access to artificial water sources (Robbins, 1993), contributes to this population-level status of good health (see section 4.1). We do note, however, that our study was conducted following a year of high rainfall (>500 mm yr^-1^ compared to 20-year average of ∼360 mm yr^-1^) and that our conclusions may differ during drought years when forage energy and protein availability is lower. Finally, wildlife act as important vectors of nutrient transport. Modifications to wildlife nutrient intake by anthropogenic mineral licks may influence the concentration and stoichiometry of nutrients across the reserve through animal-mediated lateral transport of elements, with concomitant impacts for wider ecosystem processes (Abraham et al., 2022).

In the absence of high rates of disease or predation, large herbivore population density stabilises at the ecosystem carrying capacity, which is limited by the availability of water, energy and nutrients (Chapman and Byron, 2018). As herbivore density increases, so does intra- and inter-species competition for resources, which can lead to lower body condition and resulting recruitment rates (van der Waal et al., 2003; Lane et al., 2014). Therefore, if P, Na or Zn are critically limiting for herbivore population size at TKR, could the provision of supplemental salt and mineral lick resources appreciably elevate large herbivore populations beyond the natural carrying capacity? If so, what impact could this have for conservation goals and ecosystem integrity in the long term? Over the last 20 years, TKR has experienced gradually declining veld quality due to overgrazing and drought (Tokura et al., 2018; van Rooyen and van Rooyen, 2022). Between 1999 and 2022, the mean veld condition index, measured by 111 sample plots across TKR, fell from 70% to 30%, where <40% represents low grass cover with many unpalatable annual grasses and forbs (van Rooyen and van Rooyen, 2022). Is it possible that the adopted management approach has unintentionally contributed to this degradation by decoupling wildlife health from nutrient-related feedbacks?

The southern Kalahari is a fragile ecosystem, whereby elevated herbivore population densities can have long-lasting impacts on vegetation diversity (Tokura et al., 2018). Field and remote-sensing studies at TKR have suggested that more effective predator control of large herbivore populations can help maintain vegetation productivity and diversity (Tokura et al., 2018; Makin et al., 2018). However, in the absence of sufficient predator regulation of herbivore populations in the Greater Korannaberg section, wildlife removals are conducted (∼230kg live biomass km^-2^ year^-1^), resulting in a substantial loss of nutrients from the landscape (Abraham et al., 2021). As a result, the provision of salt and mineral licks may inadvertently be generating a management conundrum, whereby licks help herbivore populations exceed the natural carrying capacity of TKR, resulting in ecological degradation and the need for off-site wildlife removal. This management process in turn leaches nutrients from the wider landscape with implications for ecosystem fertility and stoichiometry (Abraham et al., 2021), encouraging increased reliance of wildlife on mineral licks. If this hypothetical scenario is true, then the provision of anthropogenic salt and mineral licks may be misaligned from TKR’s stated goal of ecosystem restoration (https://tswalu.com/conservation-story/conservation/).

### 4.3 Management options and future research

Wildlife managers face challenges in providing supplementary nutrients for animals via free-choice salt and mineral licks. On one hand, this management strategy can rectify ethical issues related to animal husbandry. But on the other, it may elevate herbivore populations beyond the long-term carrying capacity with possible ecosystem-wide degradation (Felton et al., 2017). Each protected area will have associated idiosyncrasies, and assessments must be made on a location-by-location basis.

To ensure that provision of anthropogenic salt and mineral licks aligns with conservation goals, managers should seasonally assess nutrient levels from various resource pools including forage, water and soil. Where inter-annual variability of nutrient resources is high (e.g. due to climate change, anthropogenic disturbance or migratory/invasive species; Birnie-Gauvin et al., 2017), assessments should be undertaken more frequently. Our study provides a framework to quantify the contribution of different nutrient resource pools to wildlife diet. However, future improvements to this framework such as species-specific forage intake and camera-trap monitoring across seasons will help refine estimates. Importantly, future assessments must define what nutrient thresholds are appropriate to achieve conservation goals, balancing the needs of animal health while restoring natural processes. We note that this may be a challenge for wildlife managers as tourist often prefer to observe “healthier” animals (Dubois and Fraser, 2013), despite catabolism and death being natural processes (Robbins, 1993). For example, a rewilding experiment at Oostvaardersplassen, Netherlands, was terminated in 2018 due to public outcry over unacceptable levels of animal starvation (Theunissen, 2019). Here, we suggest that based on calculated nutrient deficiencies, only essential nutrients that correct wildlife deficiencies should be available when required. For example, at TKR, managers should consider whether licks are required during the dry season when energy and protein availability limits nutrient utilisation of large herbivore grazers (see section 4.1). This will decrease wildlife congregations at lick sites, reducing areas of high ecological degradation (Brown and Cooper, 2006; Priesmeyer et al., 2012). To ensure that nutrient intake calculations are robust, managers should regularly monitor the health of animals and adjust lick provision accordingly. Here, we have presented two non-invasive options for managers to monitor large mammalian herbivore health using faecal and body condition indicators. Other methods such as gut/tissue sampling, telemetry, stable isotopes and direct behavioural observations could also be used (Birnie-Gauvin et al., 2017).

Many protected areas are located in marginal environments (Joppa and Pfaff, 2009), which may suffer decreasing fertility over the coming century (Birnie-Gauvin et al., 2017). To reduce wildlife reliance upon mineral licks in these environments, managers should recalculate the long-term carrying capacity for their reserve (Milner et al., 2014). Lowering herbivore density will allow resident individuals to access higher quality resources via decreased intra- and inter-species competition (van der Waal et al., 2003; Gann et al., 2016; Okita-Ouma et al., 2021). Herbivore populations can be decreased by off-site removal. However, this comes with associated costs of nutrient leaching (Abraham et al., 2021), which over long periods (e.g. decades) may result in declining ecosystem fertility. Alternatively, more balanced natural predation can regulate herbivore populations, whilst increasing landscape nutrient heterogeneity (Monk and Schmidt, 2021) and creating additional nutrient resources such as bones for osteophagy (Bredin et al., 2008). Landscapes of fear generated by predators can additionally prevent prey from over-exploiting high-resource areas, including mineral lick sites (Link et al., 2010), which may further help regulate herbivore population levels and reduce ecosystem degradation associated with congregating behaviour.

In this study, we have shown that for a wildlife reserve in the Kalahari Desert, nutrients acquired from anthropogenic licks can contribute significantly to herbivore nutrient intake requirements, particularly for larger-bodied species. Simultaneously, animals appear in good health, indicating that current wildlife management practices may play a role in artificially inflating populations beyond the natural ecosystem carrying capacity in the absence of mineral licks. This mechanism has potential impacts on ecosystem integrity through overgrazing and habitat degradation. This is, however, an observational study and can only show associations, not causation. Furthermore, the conclusions drawn in our study rely heavily on research conducted on the effects of salt and mineral licks for domestic livestock and white-tailed deer (Suttle, 2010; Milner et al., 2014). Yet, nutrient resources have been supplied in wildlife landscapes across the world for hundreds, if not thousands of years (Oro et al., 2013) and are increasingly being used to actively promote conservation goals (e.g. Felton et al., 2017; Simpson et al., 2020). Further experimental investigations examining the impact of mineral lick provision on ecosystem composition and function are required. Based on results presented here, it is clear that anthropogenic provision of mineral licks should be carefully considered by wildlife managers aiming to conserve or restore natural processes in these conservation and rewilding landscapes.

## References

Abraham, A. J., Webster, A. B., Prys-Jones, T. O., le Roux, E., Smith, D., McFayden, D., … Doughty, C. E. (2021). Large predators can mitigate nutrient losses associated with off-site removal of animals from a wildlife reserve. Journal of Applied Ecology, 58(7), 1360–1369.

Abraham, A.J. et al. Understanding anthropogenic impacts on zoogeochemistry is essential for ecological restoration. Restoration Ecology.

Bartoskewitz, M. L., Hewitt, D. G., Pitts, J. S., & Bryant, F. C. (2003). Supplemental feed use by free-ranging white-tailed deer in southern Texas. Wildlife Society Bulletin, 1218–1228.

Birnie-Gauvin, K., Peiman, K. S., Raubenheimer, D., & Cooke, S. J. (2017). Nutritional physiology and ecology of wildlife in a changing world. Conservation Physiology, 5(1).

Boone, R. B., & Hobbs, N. T. (2004). Lines around fragments: effects of fencing on large herbivores. African Journal of Range and Forage Science, 21(3), 147–158.

Bredin, I. P., Skinner, J. D., & Mitchell, G. (2008). Can osteophagia provide giraffes with phosphorus and calcium? Onderstepoort Journal of Veterinary Research, 75(1), 1–9.

Brown, R. D., & Cooper, S. M. (2006). In my opinion: the nutritional, ecological, and ethical arguments against baiting and feeding white-tailed deer. Wildlife Society Bulletin, 34(2), 519–524.

Buckland, S. T., Rexstad, E. A., Marques, T. A., & Oedekoven, C. S. (2015). Distance sampling: methods and applications (Vol. 431). Springer.

Calder 3rd, W. A., & Braun, E. J. (1983). Scaling of osmotic regulation in mammals and birds. American Journal of Physiology-Regulatory, Integrative and Comparative Physiology, 244(5), R601–R606.

Chapman, E. J., & Byron, C. J. (2018). The flexible application of carrying capacity in ecology. Global Ecology and Conservation, 13, e00365.

Clauss, M., Steuer, P., Erlinghagen-Lückerath, K., Kaandorp, J., Fritz, J., Südekum, K.-H., & Hummel, J. (2015). Faecal particle size: digestive physiology meets herbivore diversity. Comparative Biochemistry and Physiology Part A: Molecular & Integrative Physiology, 179, 182–191.

Clauss, M., Steuer, P., Müller, D. W. H., Codron, D., & Hummel, J. (2013). Herbivory and body size: allometries of diet quality and gastrointestinal physiology, and implications for herbivore ecology and dinosaur gigantism. PLoS One, 8(10).

Codron, D., Codron, J., Lee-Thorp, J. A., Sponheimer, M., De Ruiter, D., Sealy, J., … Fourie, N. (2007). Diets of savanna ungulates from stable carbon isotope composition of faeces. Journal of Zoology, 273(1), 21–29.

Cox, D. T. C., & Gaston, K. J. (2018). Human–nature interactions and the consequences and drivers of provisioning wildlife. Philosophical Transactions of the Royal Society B: Biological Sciences, 373(1745), 20170092.

Cromhout, M. (2007). The ecology of the African buffalo in the Eastern Kalahari region, South Africa. University of Pretoria.

De Brouwer, C. H. M., Cilliers, J. W., Vermaak, L. M., Van der Merwe, H. J., & Groenewald, P. C. N. (2000). Phosphorus supplementation to natural pasture grazing for beef cows in the Western Highveld region of South Africa. South African Journal of Animal Science, 30(1), 43–52.

Dubois, S., & Fraser, D. (2013). A framework to evaluate wildlife feeding in research, wildlife management, tourism and recreation. Animals, 3(4), 978–994.

Ezenwa, V. O., Jolles, A. E., & O’Brien, M. P. (2009). A reliable body condition scoring technique for estimating condition in African buffalo. African Journal of Ecology, 47(4), 476–481.

Felton, A. M., Felton, A., Cromsigt, J. P. G. M., Edenius, L., Malmsten, J., & Wam, H. K. (2017). Interactions between ungulates, forests, and supplementary feeding: the role of nutritional balancing in determining outcomes. Mammal Research, 62(1), 1–7.

Gann, W. J., Fulbright, T. E., Grahmann, E. D., Hewitt, D. G., DeYoung, C. A., Wester, D. B., … Draeger, D. A. (2016). Does supplemental feeding of white-tailed deer alter response of palatable shrubs to browsing? Rangeland Ecology & Management, 69(5), 399–407.

Goff, J. P. (2018). Invited review: Mineral absorption mechanisms, mineral interactions that affect acid–base and antioxidant status, and diet considerations to improve mineral status. Journal of Dairy Science, 101(4), 2763–2813.

Grant, C. C., Peel, M. J. S., & Van Ryssen, J. B. J. (2000). Nitrogen and phosphorus concentration in faeces: an indicator of range quality as a practical adjunct to existing range evaluation methods. African Journal of Range and Forage Science, 17i1–3), 81–92.

Helary, S. F., Shaw, J. A., Brown, D., Clauss, M., & Owen-Smith, N. (2012). Black rhinoceros (Diceros bicornis) natural diets: comparing iron levels across seasons and geographical locations. Journal of Zoo and Wildlife Medicine, 43(3s).

Hobbs, N. T., & Swift, D. M. (1985). Estimates of habitat carrying capacity incorporating explicit nutritional constraints. The Journal of Wildlife Management, 814–822.

Holdø, R. M., Dudley, J. P., & McDowell, L. R. (2002). Geophagy in the African elephant in relation to availability of dietary sodium. Journal of Mammalogy, 83(3), 652–664.

Jakes, A. F., Jones, P. F., Paige, L. C., Seidler, R. G., & Huijser, M. P. (2018). A fence runs through it: A call for greater attention to the influence of fences on wildlife and ecosystems. Biological Conservation, 227, 310–318.

Joppa, L. N., & Pfaff, A. (2009). High and far: biases in the location of protected areas. PloS One, 4(12), e8273.

Kattge, J., Bönisch, G., Díaz, S., Lavorel, S., Prentice, I. C., Leadley, P., … Abedi, M. (2020). TRY plant trait database–enhanced coverage and open access. Global Change Biology, 26(1), 119–188.

Kihwele, E. S., Mchomvu, V., Owen-Smith, N., Hetem, R. S., Hutchinson, M. C., Potter, A. B., … Veldhuis, M. P. (2020). Quantifying water requirements of African ungulates through a combination of functional traits. Ecological Monographs, 90(2), e01404.

Knight, M. H., Knight-Eloff, A. K., & Bornman, J. J. (1988). The importance of borehole water and lick sites to Kalahari ungulates. Journal of Arid Environments, 15(3), 269–281.

Lane, E. P., Clauss, M., Kock, N. D., Hill, F. W., Majok, A. A., Kotze, A., & Codron, D. (2014). Body condition and ruminal morphology responses of free-ranging impala (Aepyceros melampus) to changes in diet. European Journal of Wildlife Research, 60(4), 599–612.

Leslie Jr, D. M., Bowyer, R. T., & Jenks, J. A. (2008). Facts from feces: nitrogen still measures up as a nutritional index for mammalian herbivores. The Journal of Wildlife Management, 72(6), 1420–1433.

Link, A., Galvis, N., Fleming, E., & Di Fiore, A. (2011). Patterns of mineral lick visitation by spider monkeys and howler monkeys in Amazonia: are licks perceived as risky areas? American Journal of Primatology, 73(4), 386–396.

Lintzenich, B. A., & Ward, A. M. (1997). Hay and pellet ratios: considerations in feeding ungulates. Nutrition Advisory Group Handbook Fact Sheet, 6, 1–12.

Makin, D. F., Chamaillé-Jammes, S., & Shrader, A. M. (2018). Changes in feeding behavior and patch use by herbivores in response to the introduction of a new predator. Journal of Mammalogy, 99(2), 341–350.

McDowell, L. R. (1996). Feeding minerals to cattle on pasture. Animal Feed Science and Technology, 60(3–4), 247–271.

Milner, J. M., Van Beest, F. M., Schmidt, K. T., Brook, R. K., & Storaas, T. (2014). To feed or not to feed? Evidence of the intended and unintended effects of feeding wild ungulates. The Journal of Wildlife Management, 78(8), 1322–1334.

Monk, J. D., & Schmitz, O. J. (2022). Landscapes shaped from the top down: predicting cascading predator effects on spatial biogeochemistry. Oikos, 2022(5), e08554.

Murray, M. H., Becker, D. J., Hall, R. J., & Hernandez, S. M. (2016). Wildlife health and supplemental feeding: a review and management recommendations. Biological Conservation, 204, 163–174.

Nagy, K. A., Girard, I. A., & Brown, T. K. (1999). Energetics of free-ranging mammals, reptiles, and birds. Annual Review of Nutrition, 19(1), 247–277.

O’Halloran, L. R., Shugart, H. H., Wang, L., Caylor, K. K., Ringrose, S., & Kgope, B. (2010). Nutrient limitations on aboveground grass production in four savanna types along the Kalahari Transect. Journal of Arid Environments, 74(2), 284–290.

Oesterheld, M., Sala, O. E., & McNaughton, S. J. (1992). Effect of animal husbandry on herbivore-carrying capacity at a regional scale. Nature, 356(6366), 234–236.

Okita-Ouma, B., Pettifor, R., Clauss, M., & Prins, H. H. T. (2021). Effect of high population density of eastern black rhinoceros, a mega-browser, on the quality of its diet. African Journal of Ecology, 59(4), 826–841.

Oro, D., Genovart, M., Tavecchia, G., Fowler, M. S., & Martínez-Abraín, A. (2013). Ecological and evolutionary implications of food subsidies from humans. Ecology Letters, 16(12), 1501–1514.

Pekor, A., Miller, J. R. B., Flyman, M. V, Kasiki, S., Kesch, M. K., Miller, S. M., … Lindsey, P. A. (2019). Fencing Africa’s protected areas: Costs, benefits, and management issues. Biological Conservation, 229, 67–75.

Prayag, K. D., du Toit, C. J., Cramer, M. D., & Thomson, R. L. (2020). Faunal input at host plants: Can camel thorn trees use nutrients imported by resident sociable weavers? Ecology and Evolution, 10(20), 11643–11656.

Priesmeyer, W. J., Fulbright, T. E., Grahmann, E. D., Hewitt, D. G., DeYoung, C. A., & Draeger, D. A. (2012). Does supplemental feeding of deer degrade vegetation? A literature review. In Proceedings of the Annual Conference of the Southeastern Association of Fish and Wildlife Agencies (Vol. 66, pp. 107–113).

Robbins, C. (2012). Wildlife feeding and nutrition. Elsevier.

Schultz, S. R., & Johnson, M. K. (1992). Use of artificial mineral licks by white-tailed deer in Louisiana.

Simpson, B. K., Nasaruddin, N., Traeholt, C., & Nor, S. M. (2020). Mammal Diversity at Artificial Saltlicks in Malaysia: A Targeted Use. Frontiers in Environmental Science, 8, 556877.

Skarpe, C., & Bergstrom, R. (1986). Nutrient content and digestibility of forage plants in relation to plant phenology and rainfall in the Kalahari, Botswana. Journal of Arid Environments, 11(2), 147–164.

Stapelberg, F. H., Van Rooyen, M. W., & Bothma, J. du P. (2008). Spatial and temporal variation in nitrogen and phosphorus concentrations in faeces from springbok in the Kalahari. African Journal of Wildlife Research, 38(1), 82–87.

Staver, A. C., & Hempson, G. P. (2020). Seasonal dietary changes increase the abundances of savanna herbivore species. Science Advances, 6(40), eabd2848.

Steuer, P., Südekum, K., Tütken, T., Müller, D. W. H., Kaandorp, J., Bucher, M., … Hummel, J. (2014). Does body mass convey a digestive advantage for large herbivores? Functional Ecology, 28(5), 1127–1134.

Suttle, N. F. (2010). Mineral nutrition of livestock. Cabi.

Tait, R. M., & Fisher, L. J. (1996). Variability in individual animal’s intake of minerals offered free-choice to grazing ruminants. Animal Feed Science and Technology, 62(1), 69–76.

Theiler, A., Green, H. H. & du Toit, P.J. (1924). Phosphorus in the livestock industry. Journal of the Department of Agriculture, 8(5), 460–504.

Theunissen, B. (2019). The Oostvaardersplassen Fiasco. Isis, 110(2), 341–345.

Tokura, W., Jack, S. L., Anderson, T., & Hoffman, M. T. (2018). Long-term variability in vegetation productivity in relation to rainfall, herbivory and fire in Tswalu Kalahari Reserve. Koedoe, 60(1), 1–18.

Van der Waal, C., Smit, G. N., & Grant, C. C. (2003). Faecal nitrogen as an indicator of the nutritional status of kudu in a semi-arid savanna. South African Journal of Wildlife Research, 33(1), 33–41.

Van Niekerk, B. D. H., & Jacobs, G. A. (1985). Protein, energy and phosphorus supplementation of cattle fed low-quality forage. South African Journal of Animal Science, 15(4), 133–136.

van Rooyen, N., & van Rooyen, G. (2022). Veld condition assessment of Tswalu Kalahari Reserve: Herbaceous layer.

Webster, A. B., Rossouw, R., Callealta, F. J., Bennett, N. C., & Ganswindt, A. (2021). Assessment of trace element concentrations in sediment and vegetation of mesic and arid African savannahs as indicators of ecosystem health. Science of The Total Environment, 760, 143358.

Wrench, J. M., Meissner, H. H., & Grant, C. C. (1997). Assessing diet quality of African ungulates from faecal analyses: the effect of forage quality, intake and herbivore species. Koedoe, 40(1), 125–136.

