## Supplementary Information for "Anthropogenic supply of nutrients in a wildlife reserve may compromise conservation success"

**Contents**

Supplementary text

- SI text 1: Quantification of large herbivore abundance at Tswalu Kalahari Reserve
- SI text 2: Details of water quality surveys at Tswalu Kalahari Reserve
- SI text 3: Large herbivore (>10kg) body condition assessment

Supplementary tables

- SI table 1: Large herbivore (>10kg) abundance at Tswalu Kalahari Reserve
- SI table 2: Body condition scoring categories

Supplementary figures

- SI figure 1: Estimates of TKR 2021 game counts using aerial and ground-based methods
- SI figure 2: Focal areas assessed during body condition scoring
- SI figure 3: Relative salt and mineral lick use by reserve section

References

**SUPPLEMENTARY TEXT**

**SI text 1:** Quantification of large herbivore abundance at Tswalu Kalahari Reserve

There are numerous methods to obtain estimates of animal population size with associated advantages and costs. When counting animals over large areas, such as at Tswalu Kalahari Reserve (TKR), aerial counts are often the preferred method. Aerial estimations, however, suffer from counting biases weighted to larger or more colourful animals, animals in herds or species that run from the aircraft (Jachmann, 2002). It is therefore essential that any aerial count data is validated with ground-based methods that are generally more labour intensive to collect, but do not have the same associated issues.

**Principles of distance sampling**

Here, we estimate large herbivore (>10kg) abundance at TKR using aerial and ground count data for the year 2021 using the principles of ‘distance’ sampling, the foremost technique used to estimate population sizes in marine and terrestrial ecosystems (Buckland et al., 1993). In short, a survey region is sampled by an observer systematically travelling along transects, recording any animals detected. Upon sighting an animal, the observer records the distance of the animal to the current transect. The assumptions (i) that all animals directly on a transect are detected and (ii) that detection probability decreases with distance from a transect are made. Therefore, over time, as the observer records ‘enough’ animals, a probability of detection curve can be generated. To accurately generate probability of detection curves, it is essential that (i) animal species are not misidentified, (ii) that animals do not move relative to the observer (to prevent double-counting) and (ii) that distance measurements are exact.

To transform the probability of detection curve into an estimate of animal abundance, statistical models are fitted to the raw data. These models provide an estimate of the number of animals that were statistically missed during transects. Models can range in complexity, although it should be noted that all models are only ever an approximation of the true detection fit. When smaller datasets are present, it is often useful to use expert judgement to select an appropriate model that makes logical sense (i.e. not to over-fit a model to the data). Once a model has been selected to represent the probability of detection, total animal abundance can be calculated by scaling the number of animals detected and statistically missed during transects, the total area surveyed during transects and the area of the reserve. Uncertainty estimates are generated based on how well the statistical model fits the data.

**Ground estimation (July-Aug 2021)**

To derive a ground-based estimate of large herbivore (>10kg) population sizes, we applied the ‘distance’ methodology. TKR is split into two sections separated by an electrified fence; Greater Korannaberg (92,231 ha) and Lekgaba (18,649 ha). The population size for each section was calculated independently. Ideally, transects should be straight lines placed randomly across the study region (Buckland et al., 1993). However, due to the negative ecological impact of off-road driving we utilised the pre-existing road network, which extensively covers the reserve. In many cases (e.g. along cut lines), roads were reasonably straight. However, in certain parts of the reserve (e.g. along mountain edges), transects were curved.

Recent work highlights that curved transects can still produce accurate results, provided that the observed area is reduced accordingly (Hiby and Krishna, 2001). In total, we drove 60 transects ranging between 5.7-37.5km (*μ* = 16.7, σ =9.4). To account for differences in the ways animals use space across TKR, we aimed to disperse transects across as much of the reserve as possible. Effort was made to ensure that the total transect length in each of the five main habitat types present at TKR were driven proportionally to their area (van Rooyen and van Rooyen, 2017). Additionally, all transects were conducted between 7:30am and 11am to reduce issues related to animals migrating to waterholes during the middle of the day.

During each transect, a Garmin 735XT GPS watch was used to track the route. To ensure observer consistency, the a Toyota Hilux moving at ~15km/hr driven by same driver (A. Abraham) was used throughout, and the same spotter (A. Webster) always stood elevated in the back. Upon spotting an animal, the vehicle was stopped immediately and the GPS location of the vehicle was recorded. Data pertaining to species, number of individuals, compass angle and distance from vehicle was collected. Distance from vehicle was calculated using an AOFAR HX-700N range finder. In the R statistical software v3.6.1, all data was transformed into UTM Zone 34S and minimum distance (m) of each animal observation to the driven transect was calculated using the gDistance function in the ‘rgeos’ package (Bivand and Rundel, 2019).

To account for curves in the road, we calculated the effective length driven by creating a buffer around each transect and estimating the straight-line length of the total area (Hiby and Krishna, 2001). Buffer distances were calculated for each species individually using the 99^th^ percentile minimum distance. Total transect length was 684.2km in the Greater Korannaberg section and 325.2km in the Lekgaba section. In total, the number of large herbivores (>10kg) counted in the Greater Korannaberg was 1658 individuals and 1361 individuals in Lekgaba.

For each large herbivore species (>10kg) and reserve section, we fitted three detection function models to the minimum distance data using the ‘distance’ package (Miller et al., 2019). Following Thomas et al. (2010) we used (i) half-normal key with cosine adjustments, (ii) half-normal key with Hermite polynomial adjustments and (iii) hazard-rate key with simple polynomial adjustments. These three models fit most cases and rarely are other more complex models required (Thomas et al., 2010). To generate an estimate of population size for each species, we scaled fitted models by the total area (km^2^) of each section. For species that are never or rarely observed in mountain regions, we used the section area excluding mountain regions calculated using habitat shape files generated by van Rooyen and van Rooyen (2017). Each model provides a best estimate and uncertainty based on the quality of fit to the data. We used expert judgement to discard over- fitted models. Generally, this only occurred for rare species (e.g. sable, roan, impala) where observation datasets were small. Finally, a ‘best’ population size estimate was calculated by taking the mean across remaining models. Associated uncertainty was calculated using the minimum and maximum values across models.

**Aerial estimation (March 2021)**

Aerial count data was collected at TKR between March and April 2021 using an Augusta Westland 139 helicopter flying parallel transects at a height of 100ft AGL and speed of 50-60 knots. Four observers (two port, two starboard) were used to count large herbivores (>10kg) from the helicopter. Observations are recorded at the point at which animals are perpendicular to the helicopter and classified into four distance categories (0-50m, 50-100m, 100-200m, 200-400m). Transects are spaced 800m apart to ensure that the entire area of TKR is recorded. Consequently, the data can be analysed using the same ‘distance’ methodologies outlined above for the ground transects. In order to generate probability of detection curves for the aerial data in 2021, individual animals within each distance band were randomly allocated an exact distance within that band using the ‘sample’ function in R statistical software v3.6.1. The same three detection functional models were then fitted to the data for each section and species combination using the ‘distance’ package (Miller et al., 2019). The same method was used to calculate the ‘best’ population size estimate and associated uncertainty.

**Ground and aerial population estimate comparison**

This approach facilitated a comparison between ground-based and aerial estimates for the year 2021 (SI Fig. 1). Large herbivore groups (>10kg) have been categorised based on their response to the helicopter (Jachmann, 2002). It should be noted that the aerial survey was conducted in the early dry season (late-March), whilst the ground-truthing exercise was conducted in the late dry season (July-Sept) following off-site game removal in May. Consequently, results presented here have been corrected for game removals, which affect gemsbok, eland, blue wildebeest, kudu, mountain zebra and plains zebra only.

As the helicopter approaches, some species such as steenbok, kudu and giraffe tend to freeze, whilst others such as gemsbok, wildebeest and eland tend to run away. Consequently, due to the possibility of double counting, species that flee from the helicopter transects may have consistently higher counts in the aerial methodology. In general, however, there is a very close relationship between methods (R^2^ = 0.81; SI Fig. 1) and there do not appear to be appreciable differences between estimation methods across the freeze-run continuum (SI Fig. 1).

Notable discrepancies in estimated population abundances between methodologies include:

- **Eland:** Higher estimates from aerial transects. Likely due to large herd-nature of eland, which can cause biases in model fit when data is collected over a relatively short period of time (Buckland et al., 1993). May also be influenced by animals running from helicopter (Jachmann, 2002).
- **Steenbok:** Higher estimates from ground transects. Estimates are likely underestimated from the air, as steenbok are difficult to see. It is possible that not all individuals directly on the transect are observed as detections directly beneath the helicopter are difficult to see.
- **Mountain zebra and kudu:** Higher estimates from aerial transects. These species utilise mountains for safety. Ground estimates are likely to be underestimates due to bias of road network to plains regions.

Despite the notable discrepancies outlined above, it is clear that the aerial census data has been collected rigorously and provides an accurate reflection of large herbivore (>10kg) population sizes in TKR.

**SI text 2:** Water quality surveys at Tswalu Kalahari Reserve

TKR commissioned multiple water quality assessments over the period 2000-2021. Assessments were made by African Water Solutions (October 2000), KLM Consulting Services (May 2017), KGH Applied Geological (June 2019; February 2020) and Clearwater Pumps (September 2019). Measurements were taken from surface waterholes and borehole pumped water used to supply waterholes. In total, 31 separate waterholes/boreholes were sampled (Greater Korannaberg n = 29; Lekgaba n = 2). Results for the concentration in mg litre^-1^ of sodium (Na), phosphorus (P), calcium (Ca), iron (Fe), zinc (Zn) and magnesium (Mg) were collated from individual reports. Where multiple measurements were reported for a waterhole/borehole, a mean value per waterholes was calculated. Finally, a mean value across all waterholes at TKR was calculated. Results are presented in Table 2.

**SI text 3:** Large herbivore (>10kg) body condition assessment

Examination of animal body condition can provide an external assessment of overall animal health. However, individual body condition varies throughout the year and is typically dependant on age, reproductive state (pregnant, lactating, rutting) and seasonal resource availability (Bourbonnais et al., 2016; Schiffmann et al., 2017). Despite the limitations of this approach (e.g. it is a subjective measure and difficult to quantify short-term changes in lipid content; Wilder et al., 2016), body condition scores are used extensively to assess animal fitness. The physical condition of an individual is strongly correlated to overall health (Schiffmann et al., 2017) and directly related to reproductive potential (Robbins et al., 2012), immune response (Møller et al., 1998; Sanchez et al., 2018), the ability to survive periods of nutritional stress (Verrier et al., 2011) and vulnerability to predation (Murray, 2002). For these reasons, non-invasive body condition scoring (BCS) systems are a widely accepted method for assessing the physical condition in livestock (Freitas et al., 2019). Such techniques have since been developed for and applied to specific wild ungulates in southern Africa, including buffalo (*Syncerus caffer*: Ezenwa et al., 2009), impala (Munro and Skinner, 1979) and giraffe (*Camelopardus giraffa*: Clavadetscher et al., 2021).

Different BCS assessments are available and include composite (where individual body regions are scored and a sum or mean is calculated), algorithm (where a score is achieved by following a flow chart) and overview protocols (where a score is given based on overall appearance) (Schiffmann et al., 2017). As our BSC assessment was conducted during line transect surveys, there was limited time spent with individuals. Additionally, there was uncharacteristically long grass during the survey period. Consequently, overview protocols were the most suitable approach for BC assessment in this study.

To ensure equal geographical representation and reduce bias from repeatedly measuring the same individual, body condition was evaluated during line transect assessments (see SI text 1 for details on transect design) during the dry season (July-September 2021). Upon sighting an individual/group of animals, an overall impression of BC was obtained for each individual by assessing the presence/absence of soft tissue around bony anatomical structures including the shoulders, ribs, spine, hips and tail base, (SI Fig. 2). Body condition scores between 1 (emaciated) and 5 (obese) were assigned to each individual after observation (SI Table 1). Where possible, additional factors such as coat condition (prominence of coat markings, coat condition, presence or absence of mange) and prominence of the dewlap were assessed.

For rare species, where observation during ground transects was low (n<40), additional information was obtained from camera trap data collected at mineral licks (see Section 2.4 in the main text for details on camera trap deployment). To prevent bias in this additional camera trap data, which may arise due to repeat visits by the same individual/group, we standardised the length of all camera trap deployments and randomised the order of videos from which data was obtained. In total, 1865 BCS were obtained across large herbivore (>10kg) species at TKR.

| **Species** | **Mass**  **(kg)** | **Greater Korannaberg** | | **Lekgaba** | |
| --- | --- | --- | --- | --- | --- |
|  |  | **Aerial** | **Ground** | **Aerial** | **Ground** |
| White rhino (*Ceratotherium simum*) | 2196 | - | - | - | - |
| Giraffe (*Giraffa Camelopardalis*) | 1118 | 227 ± 45 | 319 ± 99 | 28 ± 6 | 55 ± 17 |
| Black rhino (*Diceros bicornis*) | 1000 | - | - | - | - |
| Eland (*Taurotragus oryx*) | 511 | 536 ± 61 | 87 ± 62 | 80 ± 10 | 45 ± 11 |
| Buffalo (*Syncerus caffer*) | 486 | 23 ± 7 | 45 ± 44 | - | - |
| Plains zebra (*Equus quagga*) | 280 | 10 ± 4 | 4 ± 4 | 208 ± 22 | 103 ± 34 |
| Mountain zebra (*Equus zebra*) | 279 | 454 ± 58 | 17 ± 17 | 74 ± 12 | 41 ± 26 |
| Roan (*Hippotragus equinus*) | 264 | 79 ± 21 | 134 ± 63 | - | - |
| Blue wildebeest (*Connochaetes taurinus*) | 220 | 573 ± 35 | 632 ± 36 | 104 ± 21 | 82 ± 22 |
| Sable (*Hippotragus niger*) | 211 | 84 ± 13 | 125 ± 40 | - | - |
| Gemsbok (*Oryx gazelle*) | 204 | 1904 ± 44 | 1842 ± 45 | 483 ± 22 | 213 ± 23 |
| Kudu (*Tragelaphus strepsiceros*) | 202 | 409 ± 61 | 269 ± 62 | 149 ± 34 | 117 ± 35 |
| Red hartebeest (*Alcelaphus buselaphus*) | 150 | 34 ± 19 | 46 ± 20 | 126 ± 16 | 111 ± 17 |
| Warthog (*Phacochoerus africanus*) | 76 | 372 ± 52 | 466 ± 212 | 15 ± 6 | 2 ± 2 |
| Impala (*Aepyceros melampus*) | 49 | 49 ± 10 | 131 ± 104 | 272 ± 16 | 323 ± 81 |
| Springbok (*Antidorcas marsupialis*) | 35 | 172 ± 13 | 338 ± 136 | 617 ± 66 | 487 ± 67 |
| Baboon (*Papio ursinus*) | 30 | 100 | 100 | 150 | 150 |
| Common duiker (*Sylvicapra grimmia*) | 17 | 174 ± 28 | 69 ± 69 | 184 ± 48 | 47 ± 17 |
| Steenbok (*Raphicerus campestris*) | 11 | 548 ± 61 | 968 ± 62 | 294 ± 44 | 424 ± 66 |

**SUPPLEMENTARY TABLES**

**SI table 1.** Abundance estimates of large herbivore (>10kg) species at Tswalu Kalahari Reserve in 2021 using aerial and ground-based count methodologies. Ground-based estimates have been corrected for off-site wildlife removals that were conducted in May 2021, which affect gemsbok, eland, blue wildebeest, kudu, mountain zebra and plains zebra only. Baboon abundance estimated by Tswalu wildlife managers. Information pertaining to sensitive species (rhinos) has been omitted for security reasons. Some species (e.g. buffalo, roan and sable) only occur in the Greater Korannaberg section.

**SI table 2.** Body condition scoring categories for assessment of herbivore health at Tswalu Kalahari Reserve

| **Score** | **Description** | **Characteristics evaluated** |
| --- | --- | --- |
| **1** | **Emaciated** | Animal is physically weak, disorientated and emaciated.  Bony anatomical structures are prominent across the body  Complete absence of soft tissue or muscle |
| **1.5** | **Very thin** | Emaciated but not weak – complete absence of soft tissue |
| **2** | **Thin** | Animal is thin, spine still prominent and ribs visible  Very little soft tissue evident on the rump, loin and shoulder  Evidence of muscle loss in the hind quarters |
| **2.5** | **Borderline** | Hip bones, spine as well as 12^th^ and 13^th^ ribs still visible  Muscle atrophy noticeable over shoulders, loin and hindquarters  Soft tissue evident around hip bones |
| **3** | **Healthy** | Spine, hip bones and tail head are covered with soft tissue  Muscle expression in shoulder, loin and hindquarters evident  Coat is intact and markings clear |
| **3.5** | **Very healthy** | Animal has an overall smooth, sleek appearance  Hind quarters are plump and full, ribs not visible  Coat is intact and has lustre |
| **4** | **Fat** | Animal has a round, robust appearance with lustrous coat  Hind quarters and tail head are fully covered, dewlap prominent |
| **4.5** | **Very fat** | Animal has a round, robust appearance. Bone structure is no longer visible. Fat cover over the body is thick and spongy and patchiness is likely. Brisket is full. Does not generally occur in wild populations. |
| **5** | **Obese** | Bone structure is not visible. Fat encases the tailhead and mobility may be impaired by excessive fat. Does not generally occur in wild populations. |

**SUPPLEMENTARY FIGURES**

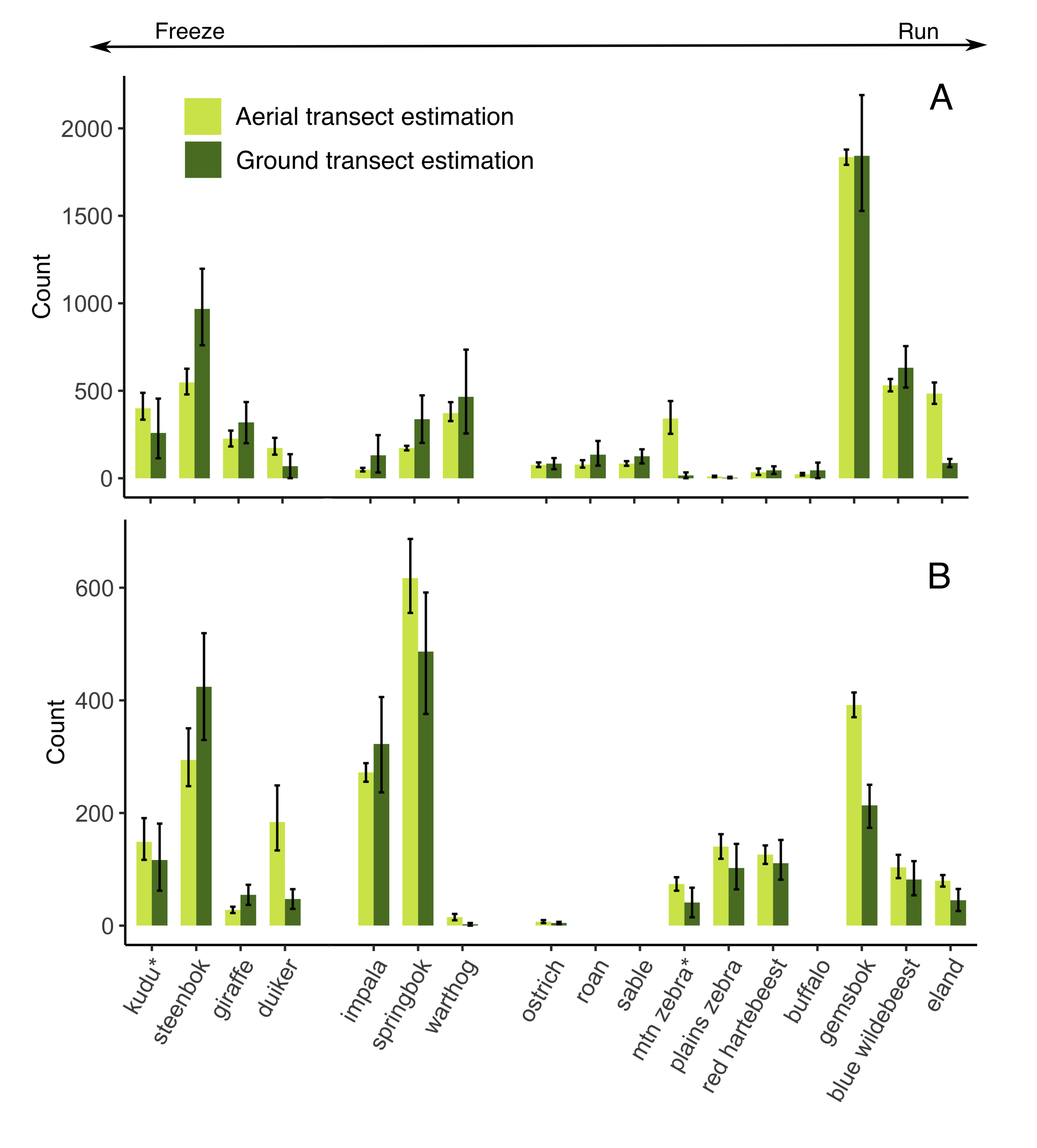

**SI figure 1:** Comparison of large herbivore (>10kg) population estimates using aerial (light green) and ground (dark green) estimate methodologies for A) the Greater Korannaberg and B) Lekgaba sections of Tswalu Kalahari Reserve in 2021. Error bars represent maximum and minimum abundance estimates from fitted probability of detection models. Species are grouped along the freeze-run continuum outlined by Jachmann (2002). Species that frequently use mountain regions are denoted by a *.

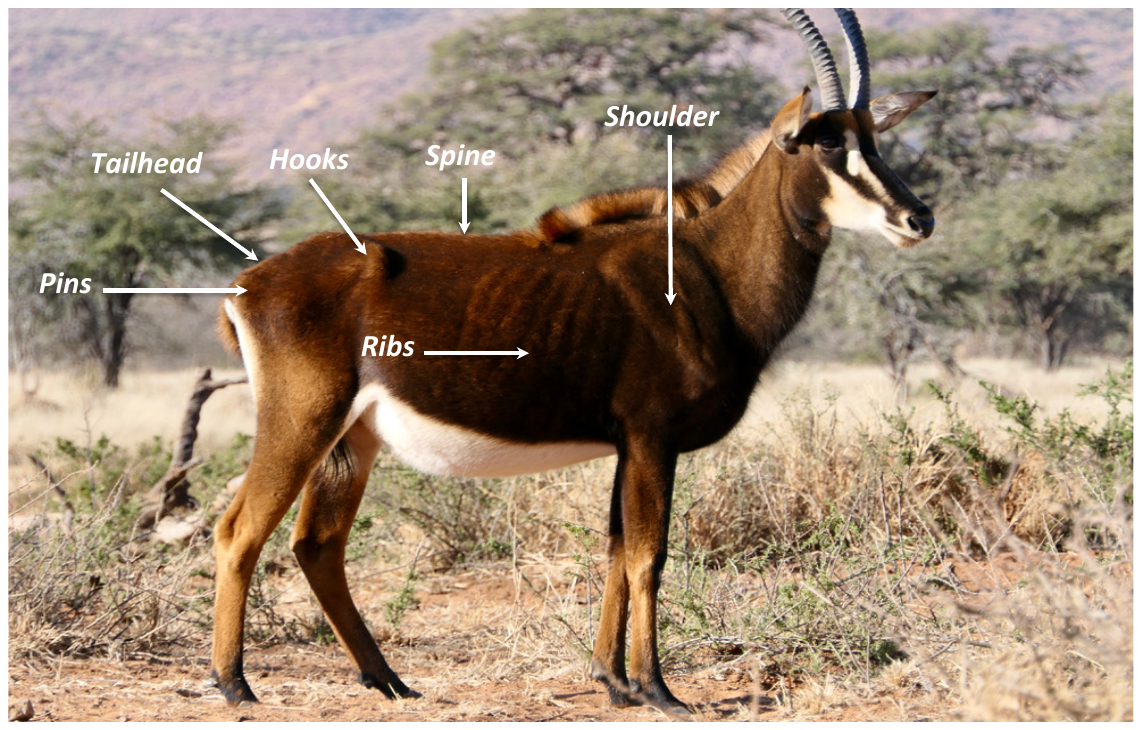

**SI figure 2.** Photograph of a female sable taken at Dedeben Pan on the Greater Korannaberg section of Tswalu Kalahari Reserve. Note the focal areas assessed during body condition scoring (BCS). Photo A. Abraham.

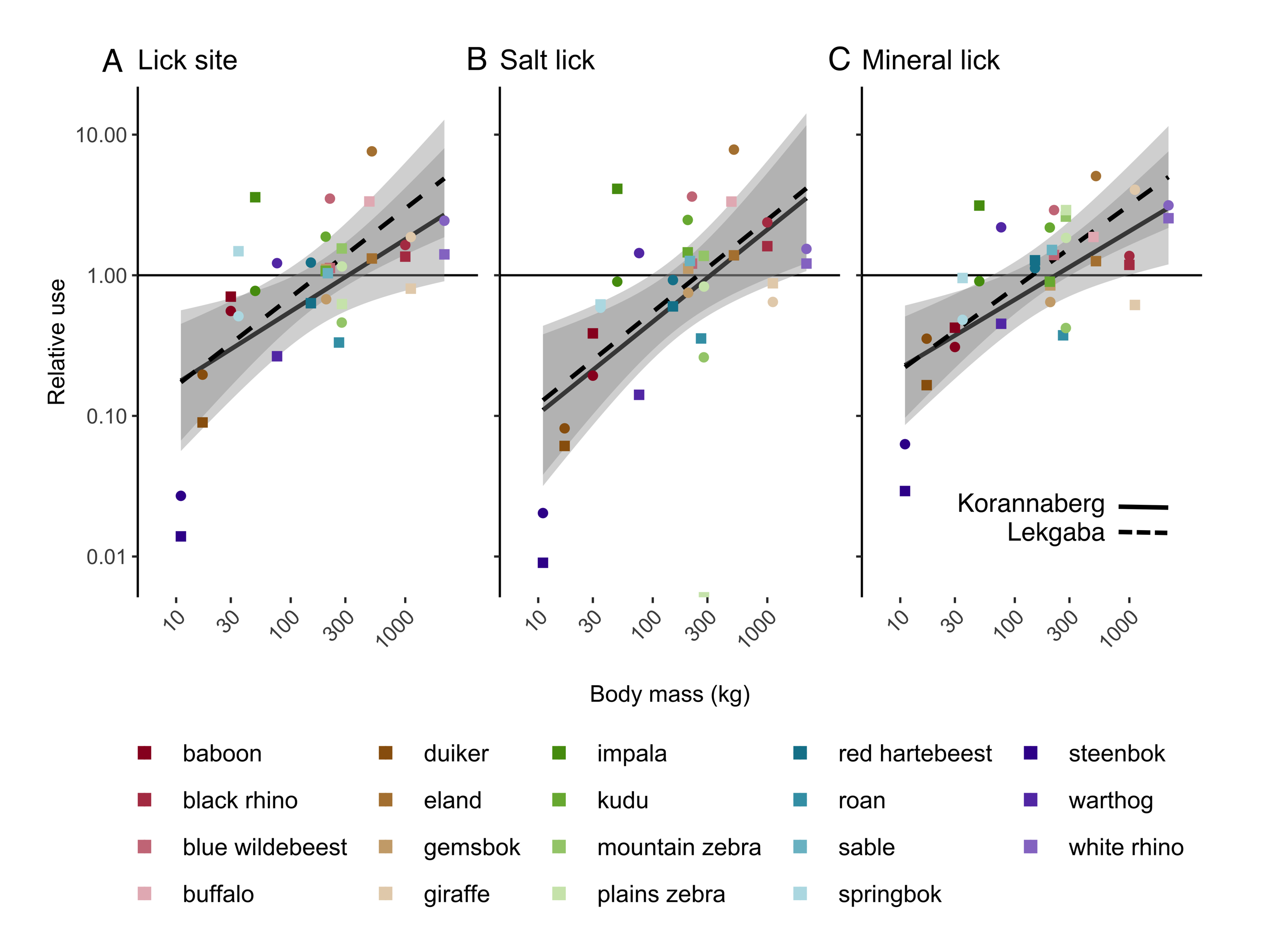

**SI figure 3.** Relative use of (A) lick site, (B) salt licks and (C) mineral licks by large herbivores at Tswalu Kalahari Reserve (TKR). Note that the y-axis is scaled log_10_, where values >1 represent more visitations of licks than expected based on species abundance and values <1 represent fewer visitations that expected. Trendlines represent a generalised-least squares model fit for all herbivores.

**REFERENCES**

Abraham AJ, Webster AB, Prys-Jones TO, le Roux E, Smith D, MacFadyen D, de Jager PC, Clauss M and Doughty (2021). Large predators can mitigate nutrient losses associated with off-site removal of animals from a wildlife reserve. Journal of Applied Ecology 2021;00:1–10.

Bivand R and Rundel (2019). Rgeos: Interface to Geometry Engine – Open Source (‘GEOS’). R Package version 0.5.2. (URL: https://CRAN.R- project.org/package=rgeos)

Bourbonnais ML, Trisalyn AN, Cattet MRL, Darimont Ct, Stenhouse GB, and Janz DM (2016). Environmental factors and habitat use influence body condition of individuals in a species at risk, the grizzly bear. Conservation Physiology 2: cou043

Buckland ST, Texstad EA, Marques TA and Oedekoven CS (2015). Distance Sampling: Methods and Applications. Springer Books

Buckland, S.T., Anderson, D.R., Burnham, K.P. and Laake, J.L. 1993. *Distance Sampling: Estimating Abundance of Biological Populations*. Chapman and Hall, London. 446pp.

Clavadetscher I, Bond MI, Martin LF, Schiffmann C, Hatt J-M and Claus M (2021). Development of an image-based body condition score for giraffes (*Giraffa camelopardalis*) and a comparison of zoo-housed and free-ranging individuals. Journal of Zoo and Aquarium Research 9: 170–185.

Ezenwa VO, Jolles AE ad O’Brien MP (2009). A reliable body condition scoring technique for estimating condition in African buffalo. African Journal of Ecology 47: 476-481.

Hiby L and Krishna MB (2001) Line transect sampling from a curving path. Biometrics 57: 727-731

Jachmann, H. (2002). Comparison of aerial counts with ground counts for large African herbivores. Journal of Applied Ecology 39: 841-852.

Miller DL, Rexstad E, Thomas L, Marshall L, Laake JL (2019). “Distance Sampling in R”. Journal of Statistical Software 89: 1-28.

Møller AP, Erritzøe CPh and Mavarez J (1998). Condition, disease and immune defence. Oikos 83: 301-306

Munro RH and Skinner JD (1979). A note on condition indices for adult male Impala (*Aepyceros melampus*). South African Journal of Animal Science 9: 47-51.

Murray DL (2002). Differential body condition and vulnerability to predation in snowshoe hares. Journal of Animal Ecology 71: 614-625.

Robbins CT, Ben-David M, Fortin JK and Nelson OL (2012). Maternal condition determines birth date and growth of newborn bear cubs. Journal of Mammalogy 93: 540-546.

Sanchez CA, Becker DJ, Teitelbaum CS, Barriga P, Brown LM, Majewska AA, Hall RJ and Altizer S (2018). On the relation between body condition and parasite infection in wildlife: a review and meta-analysis. Ecology Letters 21: 1869-1884.

Schiffmann C, Clauss M, Hoby S and Hatt J-M (2017). Visual body condition scoring in zoo animals – composite, algorithm and overview approaches in captive Asian and African elephants. Journal of Zoo and Aquarium Research 5: 1-10.

Thomas, L., Buckland, S. T., Rexstad, E. A., Laake, J. L., Strindberg, S., Hedley, S. L., ... & Burnham, K. P. (2010). Distance software: design and analysis of distance sampling surveys for estimating population size. *Journal of Applied Ecology*, *47*(1), 5-14.

Wilder SM, Raubenheimer D and Simpson SJ (2016). Moving beyond body condition indices as an estimate of fitness in ecological and evolutionary studies. Functional Ecology 30: 108-115.

van Rooyen N and van Rooyen G (2017). Ecological Evaluation of Tswalu Kalahari Reserve. An Ecological Report compiled by Ecotrust CC

Verrier D, Groscolas R, Guinet GR, Arnould JPY (2011). Development of fasting abilities in subantarctic fur seal pups: balancing the demands of growth under extreme nutritional restrictions. Functional Ecology 25: 704-717.
